# JMJD5 regulates metabolism by hydroxylating ISY1, and regulating the Arginine Methyltransferase PRMT6

**DOI:** 10.1101/2025.09.27.678987

**Authors:** Zaid A Khan, Chartinun Chutoe, Jair Marques Junior, Edward J Jarman, Andreea Gradinaru, Anthony Tumber, Eidarus Salah, Martin Lee, Haya Al Siyabi, Philippe Gautier, Chinmayi Pednekar, Ann Wheeler, Christopher J. Schofield, Luke Boulter, Alexander von Kriegsheim

**Affiliations:** Cancer Research UK Scotland Centre, Institute of Genetics and Cancer, The University of Edinburgh, Edinburgh, EH4 2XR, United Kingdom; Leicester Cancer Research Centre, University of Leicester, Leicester, LE1 5WW, UK; MRC Human Genetics Unit, Institute of Genetics and Cancer, The University of Edinburgh, Edinburgh, EH4 2XR, United Kingdom; Chemistry Research Laboratory, Department of Chemistry and the Ineos Oxford Institute for Antimicrobial Research, University of Oxford, Oxford, OX1 3TA, United Kingdom

## Abstract

2-Oxoglutarate-dependent dioxygenases (2OGDDs) employ molecular oxygen, 2-oxoglutarate, and ferrous iron to catalyse two-ectron oxidations. This dependency enables some 2OGDD to act as sensors of cellular metabolic states, driving crucial functions when oxygen or metabolic homeostasis is perturbed, including adaptation to low oxygen, epigenetic control of gene transcription, and the reshaping of metabolic pathways. Jumonji-C (JmjC) domain-containing protein 5 (JMJD5), a 2OGDD that regulates epigenetic marks, is essential for DNA damage repair and is a key regulator of cell metabolism. Notably, JMJD5 is often reduced in hepatocellular carcinoma, correlating with poor overall survival. Despite its biological significance, the molecular functions of JMJD5 remain unresolved, and its physiological targets are elusive. Here, we identify and characterise a novel signalling pathway where JMJD5 hydroxylates an arginine residue on the protein ISY1. This modification enables ISY1 to bind to and reduce the activity of Protein Arginine N-methyltransferase 6 (PRMT6). Significantly, the inactivation of PRMT6 rescues the majority of the molecular phenotype driven by JMJD5 loss, establishing the JMJD5-ISY1-PRMT6 pathway as a principal executor of JMJD5’s enzymatic function. In a genetically engineered murine liver cancer model, JMJD5 loss suppressed tumour growth and rewired one-carbon, amino acid, and lipid metabolism, recapitulating the network-level changes observed in human HCC cells. This signalling pathway clarifies existing controversies regarding JMJD5’s function and identifies PRMT6 as a potential therapeutic target for treating cancers that lack JMJD5.

## Introduction

2-Oxoglutarate-dependent dioxygenases (2OGDDs) are a diverse family of enzymes crucial for cellular metabolism, epigenetic regulation, and the response to environmental stress^1,2^. These enzymes employ molecular oxygen and 2-oxoglutarate (2OG) as cosubstrates, ferrous iron (Fe2+) as a cofactor and their activity ois often promoted by L-ascorbate, making them intrinsically sensitive to cellular metabolic and oxygen states. Amongst human 2OGDD, Jumonji-C (JmjC) domain-containing protein 5 (JMJD5), also known as KDM8, has emerged as a versatile enzyme with significant roles in both normal and pathological conditions.

In normal physiology, JMJD5 is vital for embryogenesis, circadian rhythm regulation, DNA damage repair, cell cycle control, and metabolism ^3–6^. In C. elegans JMJD5’s hydroxylase activity was shown to be required for efficient DNA damage repair, and loss of JMJD5 resulted in hypersensitivity to ionising radiation ^7^. In humans, inactivation of JMJD5 hydroxylase activity leads to severe failure to thrive, intellectual disability, and facial dysmorphism associated with increased DNA replication stress ^8^. Depletion of JMJD5 in mice leads to severe developmental defects, including embryonic lethality and growth retardation, which may be linked to the dysregulation of p53 ^9^. Taken together, these data highlight JMJD5’s role in modulating chromosome stability and metabolic pathways to maintain cellular homeostasis. JMJD5 is highly expressed in liver tissues, with its expression being nearly absent elsewhere. Given this restricted expression, JMJD5’s role in cancer is context-dependent. For instance, in cancers such as lung, breast, colon, prostate, and oral cancer, JMJD5 promotes tumour progression by inhibiting apoptosis, regulating cell cycle progression, enhancing glucose metabolism likely via the Warburg effect, and facilitating metastasis ^10–13^. Conversely, in hepatocellular carcinoma (HCC) and pancreatic ductal adenocarcinoma (PDAC), JMJD5 is reduced during tumour progression, an event associated with a poor prognosis ^14,15^. This highlights the tumour-suppressive functions of JMJD5 in tissues where it has high basal expression.

Despite the significant physiological roles of JMJD5, our mechanistic understanding of its functions is both limited and controversial. Although initially identified as a potential histone demethylase targeting H3K36me2 ^13^, recent evidence suggests this effect may be indirect and that JMJD5 may have distinct functions. While some studies propose that JMJD5 acts as an arginine-directed peptidase targeting the N-terminus of histones ^16,17^, the mechanism remains unclear. Structurally, JMJD5 clusters within the protein hydroxylase group of JmjC 2OGDDs, and there is compelling biochemical evidence that JMJD5 is indeed a protein hydroxylase. It exhibits arginine hydroxylase activity in vitro, catalysing the stereospecific C3 hydroxylation of arginyl residues in peptides from RCCD1 and RPS6 ^18^. However, despite clear activity in vitro, these hydroxylation sites have not been confirmed in cells, suggesting these proteins may not be physiological substrates.

While JMJD5 has clear physiological and pathophysiological phenotypes, insights into its role are limited by a lack of mechanistic understanding of its functions. Here, we aim to address this knowledge gap by characterising the role of JMJD5 in liver cells and tumours and elucidating the signalling network that executes these functions.

## Results

To identify pathways regulated by JMJD5 activity in liver cells, we used the hepatoblastoma cell line HepG2 as a model system, as it has highest endogenous JMJD5 expression of 22 liver cancer cell lines in the Human Protein Atlas (Extended Data Fig. 1 A). Using CRISPR/Cas9, we knocked out JMJD5 and subsequently rescued the loss by virally transducing either wild-type (WT) turboID-tagged JMJD5 or an enzymatically dead H321A mutant, which cannot chelate the Fe(II) required for catalytic activity ^13^. The expression level of the rescue constructs was equivalent to endogenous levels, although the H321A mutant consistently expressed at a higher level (Fig. 1A). Knock-out and rescue lines were validated by mass spectrometry-based proteomics detecting the loss or restoration of JMJD5 peptides (Fig. 1 A), because the available JMJD5 antibodies were non-specific in our hands.

**Figure 1:**
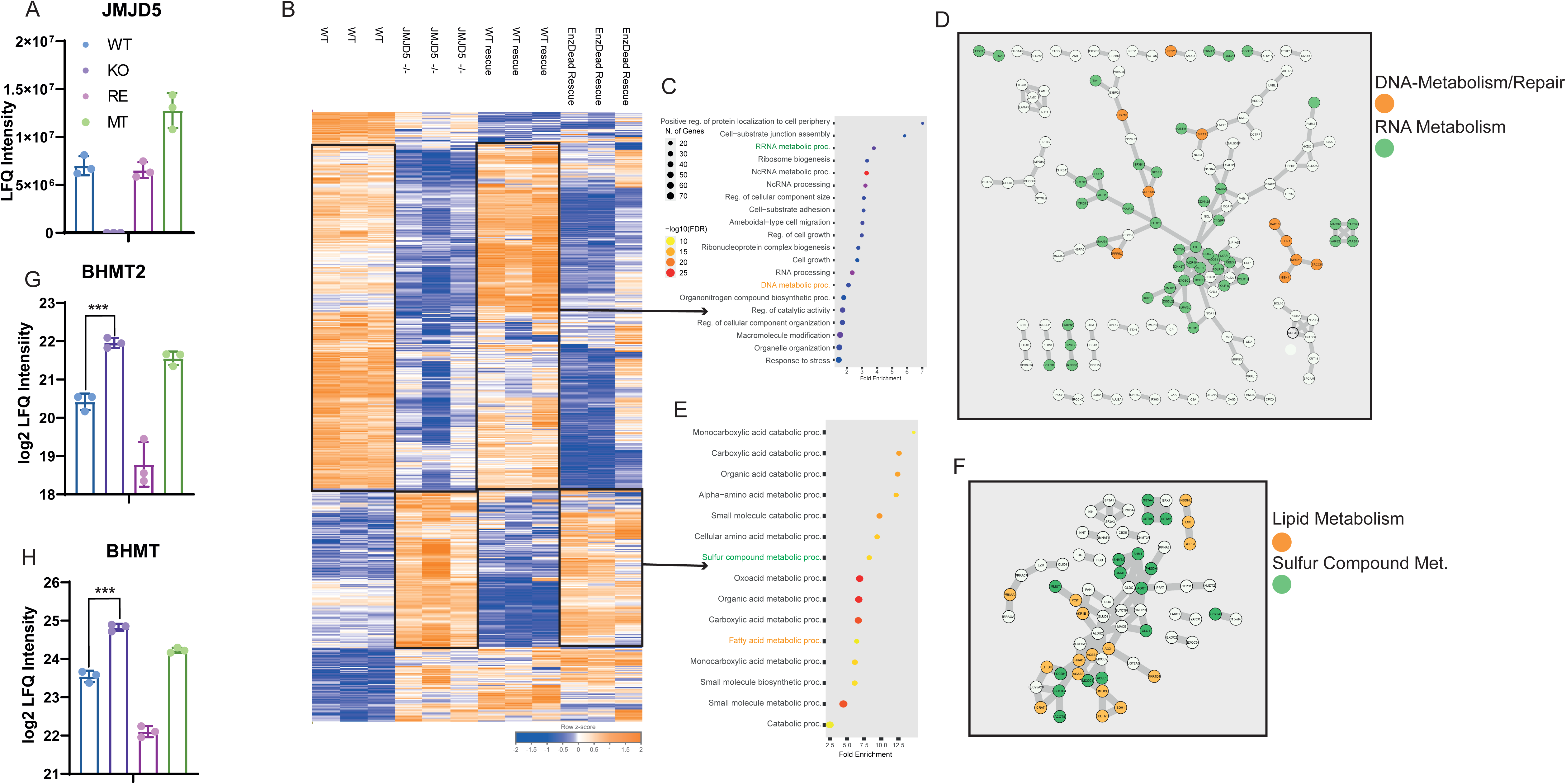
JMJD5 catalytic activity shapes the metabolic proteome of HepG2 cells. Four isogenic HepG2 lines are compared throughout: wild-type (WT), JMJD5 knock-out (JMJD5 −/−), and JMJD5 −/− cells reconstituted with either wild-type JMJD5 (RE) or the catalytically dead H321A (MT). Proteomes were measured by mass spectrometry, log2-transformed (A) Label-free quantification (LFQ) of JMJD5 protein across the four lines, confirming loss in JMJD5 −/− and re-expression of the WT (RE) and H321A (MT) constructs. Bars, mean ± s.d.; n = 3. (B) Heatmap of the proteins significantly changed between WT and JMJD5 −/−, shown for all three replicates per line and z-normalised per protein (row). Unsupervised clustering separates a cluster suppressed and a cluster induced by JMJD5 loss. (C) Gene Ontology Biological Process (GO:BP) terms over-represented in the cluster of proteins suppressed upon JMJD5 loss (Benjamini–Hochberg-adjusted p). (D) STRING protein–protein interaction network of the suppressed cluster; nodes coloured by the GO:BP categories in (C), highlighting DNA metabolism / DNA repair and RNA metabolism. (E) GO:BP terms over-represented in the cluster of proteins induced upon JMJD5 loss. (F) STRING network of the induced cluster; nodes coloured by GO:BP, highlighting lipid metabolism and sulphur-compound metabolism. (G, H) Bar graphs of log2 LFQ intensity for (G) BHMT2 and (H) BHMT — enzymes of the betaine/one-carbon arm of the methionine cycle — across the four lines (mean ± s.d., n = 3), showing induction upon JMJD5 loss and reversal on re-expression of catalytically active JMJD5. Abbreviations: LFQ, label-free quantification; GO:BP, Gene Ontology Biological Process; FDR, false-discovery rate; H321A, catalytically dead JMJD5 (MT).

### JMJD5 loss regulates metabolic networks in HepG2 cells

We then quantified the proteome of these four cell lines by mass spectrometry (Data Table 1). The loss of JMJD5 altered the expression of a surprisingly large proportion of the proteome. The majority of these changes were rescued by re-expressing WT JMJD5, whereas the H321A mutant failed to rescue the proteome-wide expression changes (Fig. 1B). This demonstrates that the enzymatic activity of JMJD5 is essential for driving the majority of its cellular functions. A subset of changes in the heatmap was not reversed by JMJD5 re-expression; we attribute this non-rescued cluster to the use of CMV-driven turboID-tagged JMJD5 rescue rather than to off-target effects, as we assayed a polyclonal knockout population where off-targets would be diluted in the pool. Closer inspection of the pathways and networks dysregulated by the loss of JMJD5 revealed that its absence reduced the expression of proteins involved in RNA metabolism, DNA metabolism, and damage repair (Fig. 1C & D, Extended Data Fig. 1B). Conversely, JMJD5 loss resulted in the induction of proteins involved in numerous metabolic pathways, including those for lipids, amino acids, and sulphur compounds (Fig. 1E & F). We then assayed whether these changes were conserved at the transcriptional level and found that JMJD5-induced proteome changes poorly correlated with the transcriptome apart from proteins involved in sulphur metabolism (Extended Data Fig. 1C & D). The latter sparked our interest, as sulphur-compound metabolism is prominently involved in one-carbon metabolism and is essential for generating methyl donors for methyltransferases. Given the proposed connection of JMJD5 with regulating histone methylation, we took a closer look at these proteins. We identified a core network of JMJD5 loss induced proteins that included BHMT and BHMT2 (Fig. 1G & H). These enzymes are essential regulators of the methionine cycle in the liver, where they regenerate methionine by catalysing the transfer of a methyl group from betaine to homocysteine (Extended Data 1 E) ^19^. In addition to upregulating BHMT/2, we detected the upregulation of other methyltransferases, such as glycine N-methyltransferase (GNMT) (Fig. 1 F). Together, these data suggested that JMJD5 may regulate the epigenetic landscape by altering one-carbon metabolism and the expression levels of key methyltransferases.

### JMJD5 interacts with ISY1 in an enzyme-activity-dependent manner

Our next aim was to investigate a mechanistic link between JMJD5’s enzymatic activity and the induction of these metabolic and methylation regulators. Since the WT, but not the enzymatically dead mutant, could rescue the observed changes in BHMT/GNMT expression, we hypothesised that the signalling downstream of JMJD5 must involve a direct substrate of JMJD5. To identify potential substrates, we designed an unbiased interaction proteomic screen based on a substrate-trap approach we previously used to successfully identify substrates of the related hydroxylase FIH ^20^. In this screen, we used the broad spectrum 2OGDD inhibitor DMOG (a prod-drug form of N-oxalylglycine) to trap the substrate-enzyme complex using the H321A mutant as a control, as its inability to chelate the catalytic iron ion likely reduces its affinity for substrates (Fig. 2 A). We transfected Flag-tagged WT or H321A JMJD5, or a vector control, into HEK293 cells in the presence of either DMSO or DMOG.

**Figure 2:**
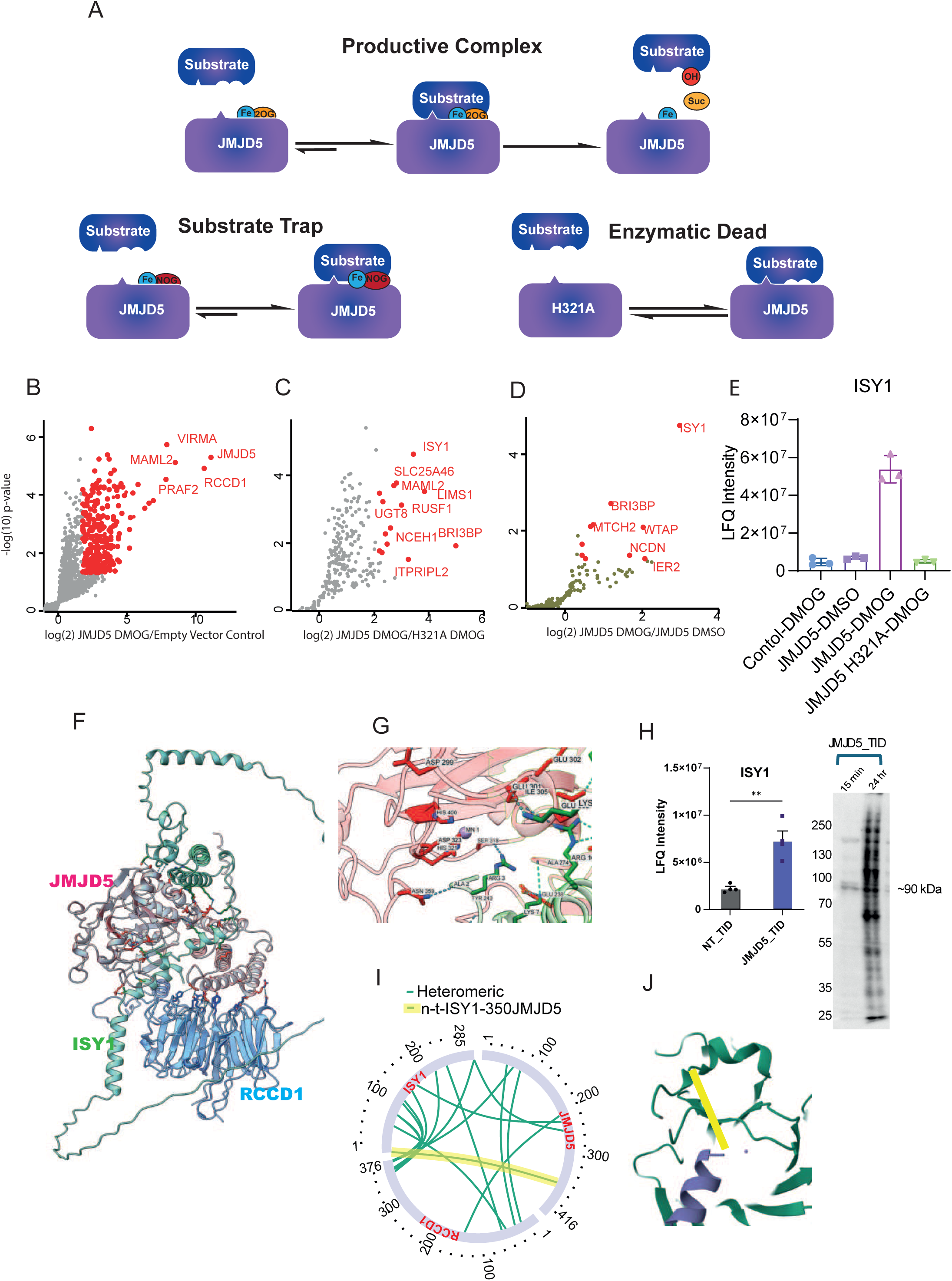
JMJD5 forms a substrate-trapping complex with ISY1. (A) Schematic of the substrate-trapping strategy. A productive enzyme–substrate complex is transient; trapping it requires either stalling catalysis with the 2-oxoglutarate analogue/inhibitor prodrug DMOG (“substrate trap”) or abolishing catalysis with the active-site mutant H321A (“enzymatic dead”), each of which stabilises the JMJD5–substrate interaction for detection. (B–D) Right-hand halves of volcano plots from JMJD5 interaction proteomics (n = 3), plotting enrichment (log2 ratio) against significance (−log10 p): (B) JMJD5 WT + DMOG vs negative control; - (C) JMJD5 WT + DMOG vs JMJD5 H321A + DMOG; - (D) JMJD5 WT + DMOG vs JMJD5 WT + DMSO (vehicle). ISY1 is selectively enriched in the catalytically stalled WT + DMOG condition. (E) Bar graph of ISY1 log2 LFQ intensity across the four interaction conditions. Bars, mean ± s.d.; n = 3. (F) AlphaFold-3 model of the trimeric JMJD5 / RCCD1 / ISY1 complex. (G) Close-up of the modelled JMJD5 active site, showing the position of ISY1 Arg3 (R3) relative to the catalytic centre. (H) Validation of proximity biotinylation. Bar graph of ISY1 log2 LFQ intensity from streptavidin pulldowns of cells expressing a TurboID-only construct (NT_TID) or a JMJD5–TurboID fusion (JMJD5_TID), n = 3, together with the corresponding streptavidin blot of the TurboID pulldown (NT_TID vs JMJD5_TID, 15 min and 24 h labelling; specific band at ∼90 kDa). (I) Cross-links detected between JMJD5, RCCD1 and ISY1 by cross-linking mass spectrometry (PhoX cross-linker; identified with xiSEARCH, visualised with xiVIEW). The ISY1 N-terminal cross-link to JMJD5 residue 350 (n-t-ISY1–350JMJD5) is highlighted in yellow. (J) The ISY1 N-terminal cross-link from (I) superimposed onto the AlphaFold structure (ISY1, purple; JMJD5, green; active-site metal, purple sphere), placing the ISY1 N-terminus at the JMJD5 active site. Abbreviations: DMOG, dimethyloxalylglycine (N-oxalylglycine prodrug); LFQ, label-free quantification; H321A, catalytically dead JMJD5.

Comparison of the co-immunoprecipitated complexes from WT JMJD5 versus the control with DMOG treatment identified the broad interactome of JMJD5 (Fig. 2 B, Data Table 2). Reassuringly, JMJD5/KDM8 itself and its best-characterised binding partner, RCCD1, were both identified as top interactors. This short-list of interactors was then cross-referenced with two additional comparisons: WT vs H321A (Fig. 2 C) and WT JMJD5 with vs without DMOG (Fig. 2 D). We focused on interactors that were enriched with DMOG treatment and showed reduced binding to the H321A JMJD5 mutant. In both comparisons, the putative splicing factor ISY1 was the most significantly enriched protein, suggesting it may be a direct JMJD5 substrate. Closer inspection revealed that the JMJD5-ISY1 interaction was only significantly induced in the presence of DMOG (Fig. 2 E). Although this interaction with JMJD5 has not been previously characterised, it was identified in a yeast-two-hybrid screen, suggesting the proteins may bind directly ^21^.

To generate a structural hypothesis for the complex, we used AlphaFold-3 ^22^ to model an interaction between JMJD5, RCCD1, which form a constitutive complex ^23^, and ISY1 (Fig. 2 F). Closer inspection of the model revealed that the N-terminus of ISY1 was positioned in close vicinity to the active site, particularly near the transition metal ion (Mn(II)) that we included in the model (Fig. 2 G). The model also predicted two hydrogen bonds between ISY1 and JMJD5 (R3-S318 and K7-E238). To validate this predicted complex, we used several orthogonal methods. Unfortunately, we failed to detect the endogenous complex, even in the presence of DMOG. However, we were able to confirm the interaction using proximity biotinylation catalysed by a turboID-JMJD5 chimaera expressed at near-endogenous levels in our JMJD5 knock-out rescue cells (Fig. 2 H and see Fig. 1 A). Furthermore, to test the prediction that the N-terminal region of ISY1 lies near the JMJD5 active site, we employed cross-linking mass spectrometry. We co-transfected RCCD1, EGFP-JMJD5, and ISY1 into HEK293 cells, immunoprecipitated the EGFP-JMJD5 complexes, and cross-linked them using the enrichable cross-linker PhoX ^24^. Analysis of the cross-linked peptides revealed one link between the N-terminal region of ISY1 and amino acid 350 in JMJD5 (Extended Data Fig. 2). Overlaying this cross-link onto our structural model showed that it was indeed compatible with the AlphaFold prediction (Figs. 2 I & J), lending further support to our model.

### JMJD5 hydroxylate Arginine 3 on ISY1

Having partially validated the structure of the JMJD5:ISY1 complex, we next devised an unbiased screen to identify post-translational modifications on ISY1 that correlate with the co-expression of active JMJD5. Again, using HEK293 cells, we expressed ISY1 in the presence of either WT or H321A JMJD5. We immunoprecipitated ISY1 and, being agnostic to the type or location of modification, used an “open” search strategy to identify all peptides derived from it ^25^. We then compared the levels of ISY1 peptides that were differentially abundant between the two conditions. Thie results revealed two peptides with statistically different levels (Fig. 3 A, Data Table 3): an N-terminal peptide (lacking the initial methionine and with N-terminal acetylation) was more abundant when co-expressed with H321A JMJD5, whereas the same peptide harbouring an additional hydroxylation on Arginine 3 (R3) was more abundant with WT JMJD5 (Fig. 3 B). Assuming roughly equal ionisation efficiencies, we estimated the occupancy of this hydroxylated site to be around 80% when WT JMJD5 was co-expressed (Fig. 3 C). Comparison of the MS/MS spectra between the hydroxylated and non-hydroxylated peptides confirmed the localisation of the +16 Da modification to R3 (Fig. 3 D). These data strongly suggest that catalytically active JMJD5 induces R3 hydroxylation on ISY1 to a high stoichiometry. JMJD5 has also been suggested to act as an arginine-directed peptidase. To investigate this, we repeated our search using a semi-specific algorithm that could identify peptides with alternative N-termini. Interestingly, we identified two such peptides, both of which were substantially induced by WT over H321A JMJD5 (Extended Figs. 3 A & B). However, the abundance of these cleaved peptides was minuscule compared to the full-length peptide, suggesting that this was either a side-reaction of JMJD5 or that the hydroxylation might induce a secondary interaction with a peptidase that becomes rate-limiting under overexpression conditions. To investigate if R3 hydroxylation is the terminal modification or if it triggers cleavage endogenously, we immunoprecipitated ISY1 from WT and JMJD5 knock-out HepG2 cells. In WT cells, we were able to identify the hydroxylated peptide but not the non-hydroxylated version. In contrast, in the JMJD5 knock-out line, we identified both forms but with reduced hydroxylation levels (Figs 3 E & F, Data Table 4). Critically, we were unable to detect any peptides consistent with N-terminal cleavage in either condition. These observations strongly suggests that ISY1 R3 hydroxylation is catalysed by JMJD5, is present on endogenous ISY1, and is the final modification. The residual hydroxylation seen in the knock-out line is likely because it is a cell pool, with approximately 5-10% of cells still expressing JMJD5 that could hydroxylate R3 during the immunoprecipitation protocol; alternatively, another hydroxylase may catalyse the reaction at a lower level. To determine whether JMDJ5 directly hydroxylated ISY1 we incubated a synthetic ISY1 N-terminal peptide with isolated recombinant JMJD5 in the presence of L-ascorbate, ferrous ammonium sulphate and 2-oxoglutarate in vitro, acquiring a time-dependent +16 Da mass shift localised to R3 (Fig. 3 G & H; Extended Data Fig. 3 C), confirming ISY1 as a direct JMJD5 substrate. Surprisingly, the efficiency of the hydroxylation was substantially increased when a 30 amino acid peptide was used over a shorter 15mer peptide (Extended Data Fig. 3D).

**Figure 3:**
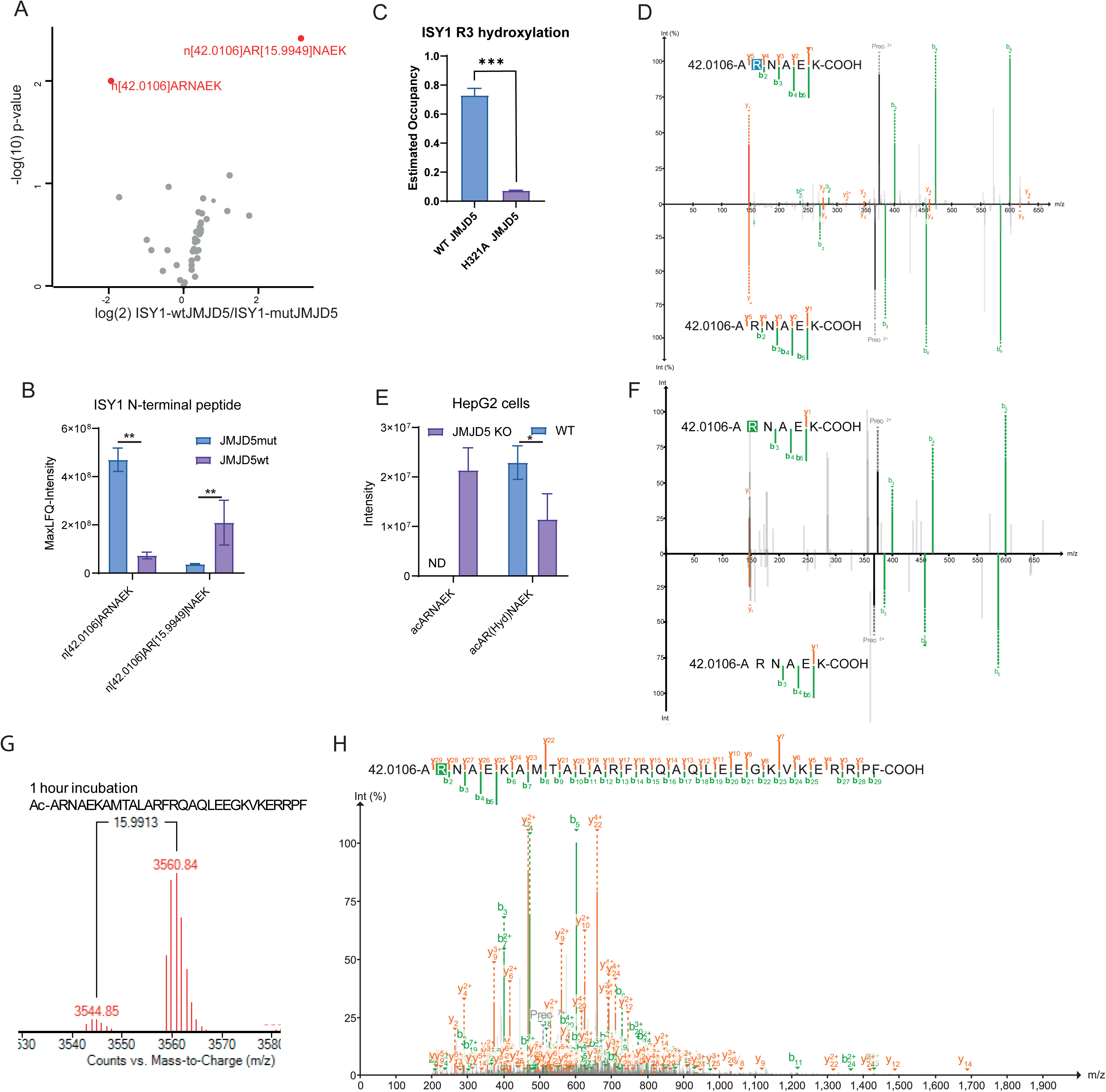
JMJD5 hydroxylates arginine 3 (R3) of ISY1. (A) Volcano plot of ISY1 peptides detected in HEK293 cells co-expressing ISY1 with either wild-type JMJD5 (JMJD5wt) or catalytically dead JMJD5 H321A (JMJD5mut), quantified with FragPipe (n = 3). The R3-hydroxylated N-terminal peptides (Acetyl-ARNAEK species) are the most strongly JMJD5wt-dependent features. (B) Bar graph of the two ISY1 N-terminal peptides differentially detected between JMJD5wt and JMJD5mut co-expression (mean ± s.d., n = 3). (C) Estimated R3-hydroxylation occupancy of ISY1 in HEK293 cells co-expressing JMJD5wt vs JMJD5mut (mean ± s.d.; p < 0.001), showing hydroxylation only with catalytically active JMJD5. (D) Representative MS/MS spectrum of the R3-hydroxylated versus non-hydroxylated ISY1 N-terminal peptide Acetyl-ARNAEK from (A); 5 ppm mass tolerance. (E) Quantification of the Acetyl-ARNAEK peptide and its R3-hydroxylated form identified on endogenous ISY1 immunoprecipitated from WT versus JMJD5 −/− (JMJD5 KO) HepG2 cells, demonstrating the modification occurs on native ISY1 in a JMJD5-dependent manner. (F) Representative MS/MS spectrum of the R3-hydroxylated versus non-hydroxylated Acetyl-ARNAEK peptide from (E); 5 ppm mass tolerance. (G) Deconvoluted mass spectrum of a synthetic ISY1 N-terminal peptide (ISY1□_□□, Ac-ARNAEKAMTALARFRQAQLEEGKVKERRPF) after 1 h in-vitro incubation with purified recombinant JMJD5, showing the +16 Da hydroxylated product. In-vitro reaction: JMJD5 (0.5 mM) with L-ascorbate (100 mM), ferrous ammonium sulphate (10 mM), 2-oxoglutarate (50 mM) and 5 mM synthetic ISY1□□□□ peptide, 1 h at room temperature. (H) MS/MS fragmentation localising the +16 Da modification specifically to ISY1 Arg3 (R3). Abbreviations: JMJD5wt/JMJD5mut, wild-type / H321A catalytically dead JMJD5; KO, knock-out; Ac, N-terminal acetyl; 2-OG, 2-oxoglutarate.

### Hydroxylation of ISY1 induces an interaction with PRMT6

Hydroxylation sites on amino acid side chains often alter protein-protein interactions, as famously exemplified by the hydroxylation-dependent binding of HIF1α to VHL ^26,27^. We therefore hypothesised that the hydroxylation of R3 on ISY1 may create or destroy a protein-binding site. To identify such hydroxylation-dependent interactions, we devised another interaction-proteomic screen. We transfected HEK293 cells with either WT ISY1 (in the presence of WT or H321A JMJD5 to modulate hydroxylation) or with ISY1 point mutants (R3A, R3K) that cannot be hydroxylated. The presence or absence of hydroxylation on R3 did not fundamentally alter the overall interactome of ISY1, with most interactions retained regardless of hydroxylation status (Figs 4 A & B, Data Table 5). To pinpoint differential binders, we generated a short list of high-confidence ISY1 interactors and used an ANOVA test to find proteins whose binding varied significantly across our conditions (Fig. 4 C).

**Figure 4:**
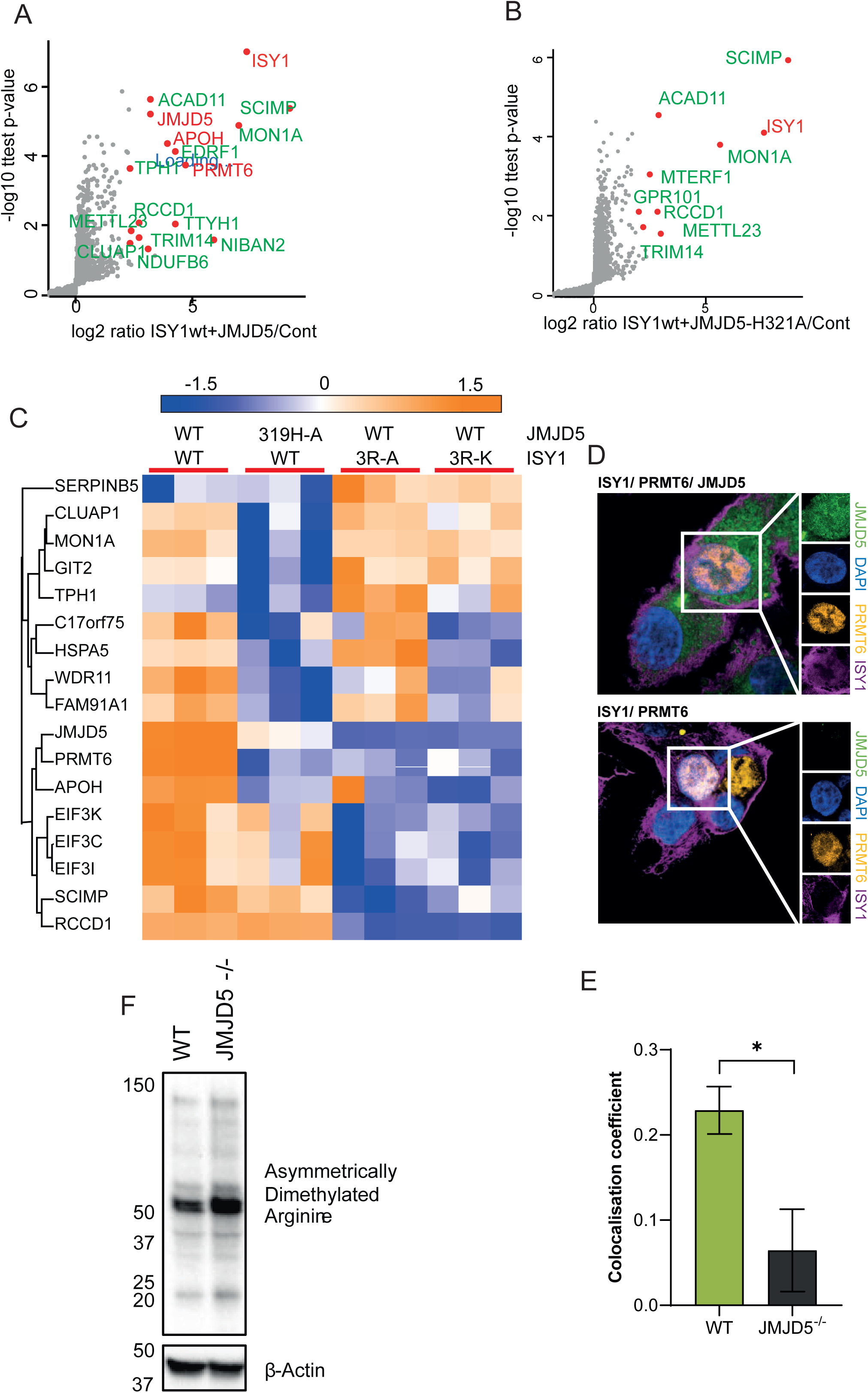
R3 hydroxylation drives formation of a PRMT6–ISY1 complex and nuclear relocalisation of ISY1. (A) ISY1 interactome in HEK293 cells: right-hand half of the volcano plot of proteins co-purifying with ISY1 when co-expressed with wild-type JMJD5, relative to a negative control co-expressing JMJD5 (n = 3). PRMT6 is enriched among ISY1 interactors. (B) As in (A) but with ISY1 co-expressed with catalytically dead JMJD5 H321A (n = 3); the OH-R3-dependent PRMT6 enrichment is lost. (C) Heatmap of ISY1-interacting proteins whose binding changes significantly across the indicated conditions (WT, H321A, R3A and R3K backgrounds), n = 3, ANOVA with permutation FDR q < 0.05, z-normalised. PRMT6 binding tracks with R3 hydroxylation. (D) Super-resolution microscopy of the reconstituted JMJD5/PRMT6 double-knockout (dKO) HepG2 lines: JP cells, which re-express eGFP-JMJD5 together with V5-mKate2-PRMT6, and P cells, which express V5-mKate2-PRMT6 only (no JMJD5). Single channels (eGFP-JMJD5 [GFP-Booster], mKate2-PRMT6 [intrinsic fluorescence], endogenous ISY1 [immunofluorescence, Alexa Fluor 647] and DAPI nuclear counterstain), merge and magnified insets are shown. PRMT6 is near-exclusively nuclear in both lines; ISY1 relocalises to the nucleus and partially co-localises with PRMT6 in JP cells (JMJD5 present), but remains cytoplasmic in P cells (JMJD5 absent), demonstrating that ISY1 nuclear localisation and PRMT6 association are JMJD5-dependent. (E) Quantification of ISY1–PRMT6 co-localisation comparing JP (JMJD5-replete; plotted as “WT”) versus P (JMJD5-deficient; plotted as “JMJD5 −/−”) cells (mean ± s.d.; p < 0.05). (F) Western blot of WT and JMJD5 −/− HepG2 lysates probed with an antibody to asymmetric dimethyl-arginine (ADMA) and β-actin (loading control), showing increased global ADMA upon JMJD5 loss.

Reassuringly, this analysis confirmed that ISY1 bound to JMJD5/KDM8 and that this interaction was reduced with the H321A mutant. Importantly, mutation of R3 to either A or K completely ablated the binding between ISY1 and JMJD5 (Fig. 4 C), indicating that this specific residue is essential for the interaction and indirectly confirming that R3 is the major hydroxylation site. We then focused on proteins that bound ISY1 in a hydroxylation-dependent manner. We were unable to detect any proteins that bound specifically to non-hydroxylated ISY1. However, two proteins, APOH and PRMT6, co-immunoprecipitated with ISY1 only when it was hydroxylated (Fig. 4 C). APOH (apolipoprotein H) is predominantly a plasma protein and an unlikely physiological binder for the nuclear protein ISY1, suggesting it may be a spurious interaction. In contrast, Protein Arginine N-methyltransferase 6 (PRMT6) is a predominantly nuclear protein and a known regulator of the epigenetic landscape ^28–30^. Moreover, PRMT6 has been shown to drive hepatic lipogenesis and alter metabolism as well are regulate DNA-damage repair ^31–33^, a phenotype reminiscent of pathways that we detected to be impacted upon JMJD5 loss. To localise these proteins, we performed super-resolution microscopy of HepG2 JMJD5/PRMT6 knockout lines that were rescued by re-expressing mKATE2-PRMT6 as well as GFP-JMJD5 or not (Fig. 4 D & E). JMJD5 was both nuclear and cytoplasmic, PRMT6 was near-exclusively nuclear, and ISY1 was predominantly cytoplasmic. In the absence of JMJD5, ISY1 remained cytoplasmic, whereas re-expression of JMJD5 permitted nuclear ISY1, and a faint but reproducible ISY1/PRMT6 nuclear co-localisation was visible (Fig. 4 E). These localisations are consistent with the JMJD5 hydroxylating ISY1, which then binds to PRMT6.

To explore the structural basis of this interaction, we generated an AlphaFold model of the PRMT6-ISY1 complex (Extended Data 4 A). The model predicted a close association of the N-terminus of ISY1 with the C-terminal domain of PRMT6 (Extended Data Fig. 4 B), regardless of hydroxylation status. However, when mapping intermolecular hydrogen bonds, the model suggested that the hydroxylation of R3 induces a new hydrogen bond between the N-terminus of ISY1 and PRMT6, potentially explaining why the modification is required for the interaction (Extended Data Fig. 4 C). We attempted to validate this complex using cross-linking mass spectrometry but were unable to identify credible linked peptides, possibly due to a weak or transient interaction.

PRMT6 predominantly catalyses the asymmetric dimethylation of arginine (ADMA) residues in proteins. To test whether JMJD5 loss might alter PRMT6 activity, we compared global ADMA levels in WT and JMJD5 knock-out HepG2 cells. We observed that ADMA levels were broadly upregulated in the knock-out cell line (Fig. 4 F), suggesting that the loss of JMJD5 leads to a gain-of-function or disinhibition of PRMT6 activity.

### JMJD5 and PRMT6 are components of a novel signalling pathway

To determine whether ISY1 and PRMT6 are essential effectors downstream of JMJD5, we aimed to knock them out in both the WT and JMJD5 knock-out cell lines. Loss of ISY1 was lethal in HepG2 cells, consistent with its classification as a common essential gene in the DepMap database ^34,35^, precluding further study. PRMT6, on the other hand, was readily knocked out by CRISPR. We hypothesised that if PRMT6 is a key downstream effector of JMJD5, then its loss should either rescue or phenocopy the signalling events caused by JMJD5 loss.

To test this, we generated two additional cell lines by knocking out PRMT6 in both WT and JMJD5 knock-out HepG2 cells (Extended Data Fig. 5 A) and analysed the proteomes of all four lines. We first identified proteins whose expression was significantly regulated by the loss of JMJD5 and then assessed how the additional loss of PRMT6 impacted their expression (Fig. 5 A, Data Table 6). While loss of PRMT6 in the WT background had almost no overlap with the JMJD5-loss signature, its ablation in the JMJD5 knock-out background rescued around half of the expression changes induced by JMJD5 loss. We isolated two clusters of proteins: those induced by JMJD5 loss and rescued by PRMT6 loss, and those suppressed by JMJD5 loss and rescued by PRMT6 loss. Gene Ontology analysis of the suppressed-rescued cluster revealed high enrichment of proteins regulating RNA metabolism and DNA repair (Fig. 5 B), while the induced-rescued cluster was enriched for sulphur amino acid/compound and lipid metabolism (Fig. 5 C). This demonstrates that these expression changes are dependent on the JMJD5-PRMT6 pathway. Closer inspection revealed that BHMT and BHMT2, key nodes in liver-specific one-carbon metabolism, were among the proteins whose induction was rescued by PRMT6 loss (Fig. 5 D).

**Figure 5:**
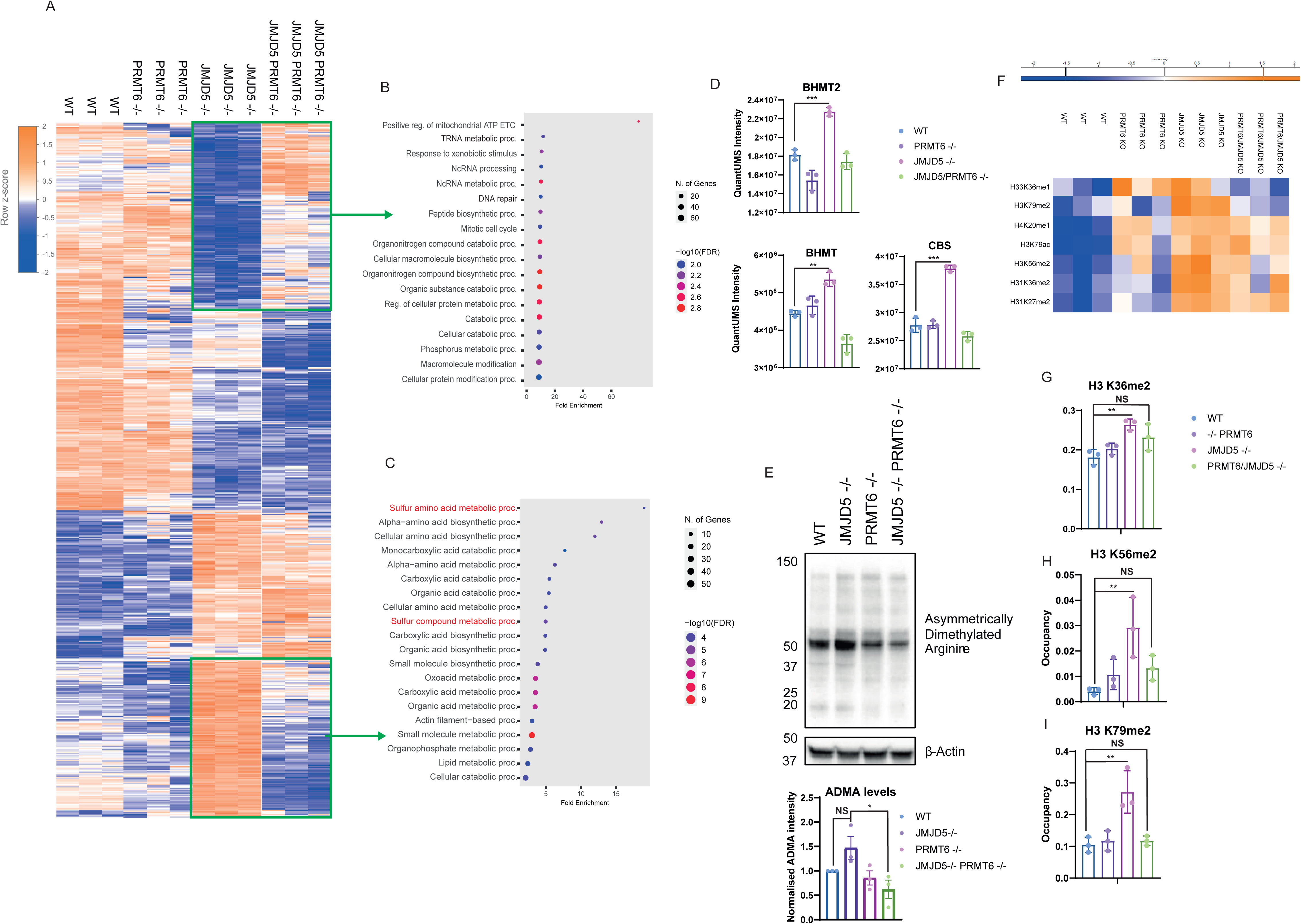
PRMT6 is a major downstream effector of JMJD5 function. Four isogenic HepG2 lines are compared: WT, JMJD5 −/−, PRMT6 −/−, and the JMJD5/PRMT6 double knock-out (DKO). Proteomes were quantified by mass spectrometry and analysed in Perseus. (A) Heatmap of proteins significantly changed between WT and JMJD5 −/− (z-normalised per protein, triplicates). Clusters outlined in green are those in which loss of PRMT6 (in the DKO) reverts the JMJD5-driven change toward WT. (B) GO:BP terms enriched in the protein cluster reduced by JMJD5 loss and rescued by concurrent PRMT6 knock-out. (C) GO:BP terms enriched in the protein cluster induced by JMJD5 loss and rescued by concurrent PRMT6 knock-out. (D) QuantUMS protein intensities of three methionine/one-carbon-cycle enzymes across the four lines (mean ± s.d., n = 3 (E) Western blot of the four lines probed for asymmetric dimethyl-arginine (ADMA) and β-actin. (3-replicate quantification; loading-corrected). (F) Heatmap of global histone post-translational modifications (PTMs) significantly regulated by JMJD5 loss across the four lines (EpiProfile quantification; n = 3; q < 0.05; z-normalised). (G–I) Bar graphs of occupancy for selected histone-PTM sites across the four genotypes (EpiProfile; pairwise t-tests; y-axes start at zero). Of the histone marks significantly altered by JMJD5 loss

To test directly whether a related methyltransferase, PRMT1, which is thought of as the dominant cellular ADMA writer, could rescue the JMJD5-loss phenotype, we generated PRMT1 KO and a JMJD5/PRMT1 double-KO HepoG2 cell lines and profiled the proteomes of all genotypes (Data Table 7, Extended Data Fig. 5 B & C). Despite only being able to achieve a partial knockout, the knockout of PRMT1 had a dramatic effect on the cellular proteome (Extended Data Fig. 5 D), affecting the proteome to a much larger extent than either JMJD5 or PRMT1 knockout. Despite this, loss of PRMT1 was unable to rescue a large proportion of the proteome changes induced by JMJD5 knockout. In some protein clusters, loss of PRMT1 compensated for the loss of JMJD5, but the effect appeared additive, suggesting an overlap or parallel pathway rather than a direct dependency. Crucially, whereas PRMT6 loss rescued expression of BHMT/2 and CBS (Fig. 5 D), loss of PRMT1 led to an augmentation of BHMT/2 expression and an additive rescue of CBS (Extended Data Fig. 5 E), overall suggesting that PRMT1 is not a downstream effector of JMJD5, and may act independently.

To confirm that the increase in ADMA upon JMJD5 loss was driven by PRMT6, we performed Western blotting on the cell lysates (Fig. 5 E). As seen before, loss of JMJD5 induced ADMA, and loss of PRMT6 alone marginally reduced ADMA levels in WT cells. Crucially, knocking out PRMT6 in the JMJD5-deficient background completely rescued the ADMA induction, demonstrating that PRMT6 is an effector of ADMA methylation driven by JMJD5 loss. In contrast, reduction of PRMT1 expression did not rescue JMJD5-driven induction of ADMA levels in the double-knockout line, again suggesting that these two proteins are not dependent on each other (Extended Data Fig. 5 F). Finally, we addressed the long-standing observation that JMJD5 loss induces H3K36me2. This histone mark induction is a consistent phenotype of JMJD5 loss across multiple systems ^3,7,13^, though it may occur via an indirect effect ^18^. We sought to determine if we could recapitulate this in HepG2 cells and whether it was dependent on PRMT6. Using a protocol for unbiased profiling of histone PTMs ^36^, we found that several H3 PTMs were indeed induced by JMJD5 loss, including methylation on K27, K36, K56, and K79. Significantly, the simultaneous loss of PRMT6 was able to at least partially rescue some of these marks, including H3K79me2, H3K56me2, and H3K36me2 (Fig. 5 F-I). Overall, these data demonstrate that a significant proportion of the signalling downstream of JMJD5 is dependent on PRMT6, establishing it as an effector of the pathway.

### JMJD5 loss regulates liver tumour growth in a genetically engineered murine model

Having established that JMJD5 loss impacts several metabolic pathways frequently dysregulated in cancer, we next wanted to determine whether loss of JMJD5 affects hepatic tumour growth in vivo. JMJD5 has a complex, context-dependent role, acting as either a tumour suppressor or promoter. To determine whether JMJD5 or RCCD1 expression is regulated in HCC we analysed normal liver tissue, tumour-adjacent and tumour tissue in transcriptomics datasets from the GTEx and TCGA consortia, respectively. In accordance with previous reports ^14^, JMJD5 mRNA was diminished in the tumour tissue, whereas RCCD1 expression was induced (Extended Data Fig. 6 A & B). Given the large discordance we observed between the transcriptome and proteome when driven by JMJD5 in cells, we wanted to determine whether protein expression levels followed a similar trend. We mined the CPTAC HBV-related HCC proteogenomic cohort ^37^ and found that JMJD5 is indeed downregulated in HCC compared to matched adjacent tissue (Extended Data Fig. 6 C). In addition, the constitutive interactor of JMJD5, RCCD1, is also downregulated at the protein but not transcript level, indicating a significant post-translational component consistent with the suggested co-stabilisation of both proteins when present in a complex (Extended Data Fig. 6 D).

Having established that JMJD5 is reduced in HCC we wanted to determine the relevant context for human HCC. Consequently, we interrogated the LIHC TCGA database and found that tumours with low JMJD5 expression frequently had mutations in β-catenin and p53 (Extended Data Fig. 6 E). The link to p53 is plausible, as JMJD5 is required for efficient DNA damage repair. Its loss in mice induces p21 and is embryonically lethal, suggesting that p53 ablation may be necessary to tolerate JMJD5 loss. Therefore, to recapitulate this common mutational background, we used a mouse model where p53 was ablated and truncated active CTNNB1 (N90 β-catenin). We also expressed c-Myc ectopically (Fig. 6 A), as the models did not develop tumours when only p53 and CTNNB1 were modified in the timeline we investigated (Data not shown). Immunohistochemistry (IHC) confirmed successful oncogene delivery (p53, MYC, beta-catenin) and the expected downstream changes in tumour sections (Extended Data Fig. 6 F). JMJD5 IHC did not yield a specific signal with the available antibodies, so the knock-out was at the bulk proteome level by mass spectrometry (Extended Data Fig. 6 G).

**Figure 6:**
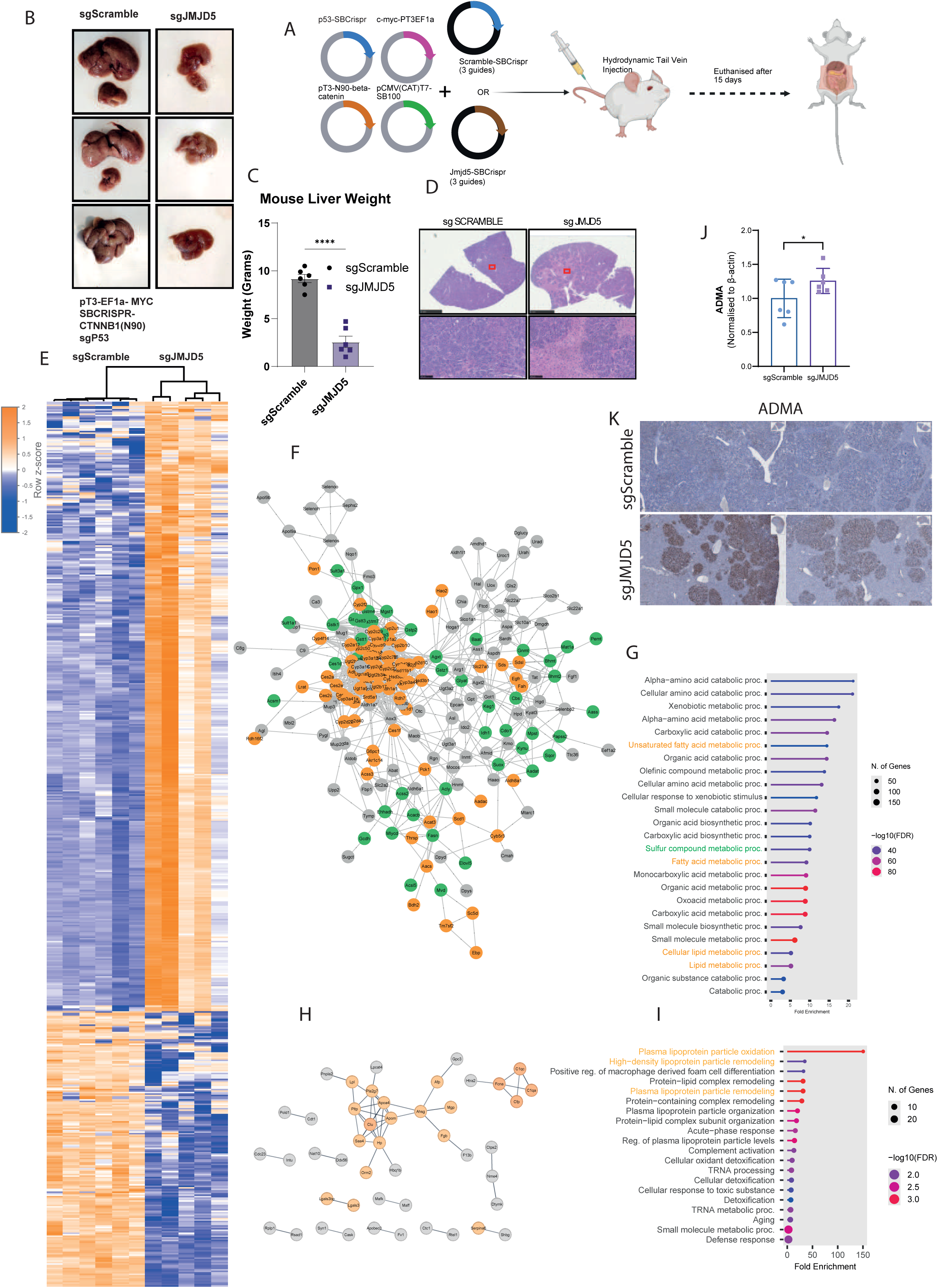
JMJD5 loss alters liver-tumour growth and signalling in vivo. A hydrodynamic tail-vein injection (HTVI) mouse liver-tumour model was used, comparing animals receiving a scrambled control gRNA (sgScramble) with a JMJD5-targeting gRNA (sgJMJD5). (A) Schematic of the experimental design (oncogene/CRISPR delivery by HTVI, tumour induction and end-point analysis). (B) Representative photographs of dissected livers from the indicated genotypes. (C) Liver weight expressed as a percentage of total body weight, by group (mean ± s.d.; n = 6 per group). JMJD5 loss reduces tumour burden. (D) Representative haematoxylin & eosin (H&E) histology; lower panel is a magnification of the boxed region. (E) Heatmap of proteins significantly changed between sgScramble and sgJMJD5 livers (q < 0.05; n = 6 and 5; z-normalised). (F) STRING network of the protein cluster induced in sgJMJD5 livers; nodes coloured by the GO:BP categories in (G). (G) GO:BP terms enriched in the induced cluster, highlighting lipid and sulphur metabolism. (H) STRING network of the protein cluster suppressed in sgJMJD5 livers; nodes coloured by the GO:BP categories in (I). (I) GO:BP terms enriched in the suppressed cluster, highlighting secreted lipoprotein/plasma-lipoprotein-particle transport proteins. (J) Violin plot of liver ADMA levels quantified by Western blotting (n = 6 and 5; 3 technical replicates per animal), showing increased ADMA on JMJD5 loss. (K) ADMA immunohistochemistry of tumour sections (sgScramble vs sgJMJD5

Loss of JMJD5 significantly reduced the size and tumour burden of the liver (Figs 6 B & C). In the livers expressing JMJD5, the majority of the tissue appeared cancerous, whereas the knock-out of JMJD5 reduced the tumour burden of the tumour, with a significant proportion of cells resembling untransformed hepatocytes (Fig. 6 D).

Histological analysis of liver tissue from mice infected with sgScramble or sgJMJD5 reveals distinct differences in tumour characteristics and cellular features. In the sgScramble group, liver tissues developed well-differentiated hepatocellular carcinoma exhibiting a solid tumour phenotype. In contrast, the sgJMJD5 group displayed a trabecular phenotype with reduced tumour invasion. Additionally, multinucleated cells were prominent in the sgScramble group, while the sgJMJD5 group presented signs of karyorrhexis in the hepatic parenchyma. In addition, we found that sgJMJD5 presented an increased sign of unresolved DNA double-strand breaks as manifested by an increase in γ-H2AX (Extended Data Figure 6 H). Macroscopically, clonal nodules were distributed across the liver in both groups, but in the JMJD5 KO livers, they appeared smaller and more restricted to the periphery, consistent with reduced expansion, possibly as a consequence of the elevated DNA-damage burden following JMJD5 loss. To determine which molecular signalling pathways were dependent on JMJD5 in this murine HCC model, we dissected the HCC tissue and analysed it by proteomics (Data Table 8). Differential expression analysis revealed two clusters of proteins or regulated by JMJD5 expression (Fig. 6 E). Similar to our observations in HepG2 cells, proteins induced upon JMJD5 loss in our in vivo model included those regulating lipid and sulphur compound metabolism (Figs 6 F & G). Conversely, proteins associated with plasma lipoprotein and lipid transport were significantly downregulated (Figs 6 H & I). Overall, at the network level, we observed a high degree of consistency for JMJD5-dependent molecular pathways between HepG2 cells and our genetically engineered murine HCC model. Finally, to determine whether ADMA was regulated by JMJD5 loss in vivo, we quantified it by Western blotting and observed that loss of JMJD5 significantly induced ADMA (Fig. 6 J, Extended Data Figure 6 I). To determine whether the increase in ADMA was localised to the tumour cells, we visualised ADMA by IHC in liver sections. We detected a dramatic increase in ADMA levels in sgJMJD5, specifically localised to the tumour section (Fig. 6 K). The IHC data recapitulated the Western blot data, demonstrating a surprisingly robust increase in ADMA when comparing sg JMJD5 to sgScramble tumours.

### JMJD5 loss in liver tumour disrupts one-carbon metabolism, redox balance, amino acids metabolism and lipid homeostasis

To investigate the metabolic consequences of JMJD5 deficiency in the liver tumour tissues, we integrated proteomics (Data Table 8), metabolomics (Data Table 9) and lipidomics data (Data Table 10). This combined analysis revealed alterations in redox regulation, one-carbon metabolism, amino acid utilisation, urea cycle and lipid remodelling.

Joint pathway analysis of up-regulated significant proteins and metabolite changes identified several significantly dysregulated metabolic pathways, including retinol metabolism, linoleic acid metabolism, glutathione metabolism, steroid biosynthesis, and cysteine and methionine metabolism (Fig. 7 A). Retinol metabolism enrichment was supported by the coordinated upregulation of multiple retinol and retinal-processing enzymes, including Rdh7, Rdh16f2, Sardh, and Adh4 (Fig. 7 B). Adh4 catalyses the oxidation of retinol to retinal, providing substrate for retinoic acid synthesis ^38^, a potent regulator of hepatic lipid metabolism and RXR-mediated transcriptional programs that influence peroxisomal function ^39^. The concurrent elevation of Rdh7 and Rdh16f2 further supports enhanced retinoid turnover, potentially contributing to the transcriptional changes and lipid remodelling observed in JMJD5-deficient livers. In parallel, proteomic analysis revealed broad activation of the glutathione S-transferase (GST) family, with 10 isoforms significantly up-regulated in JMJD5-deficient livers (Fig. 7 C). GSTs are canonical phase II detoxification enzymes with established roles in carcinogen metabolism and oxidative stress defence ^40^.

**Figure 7:**
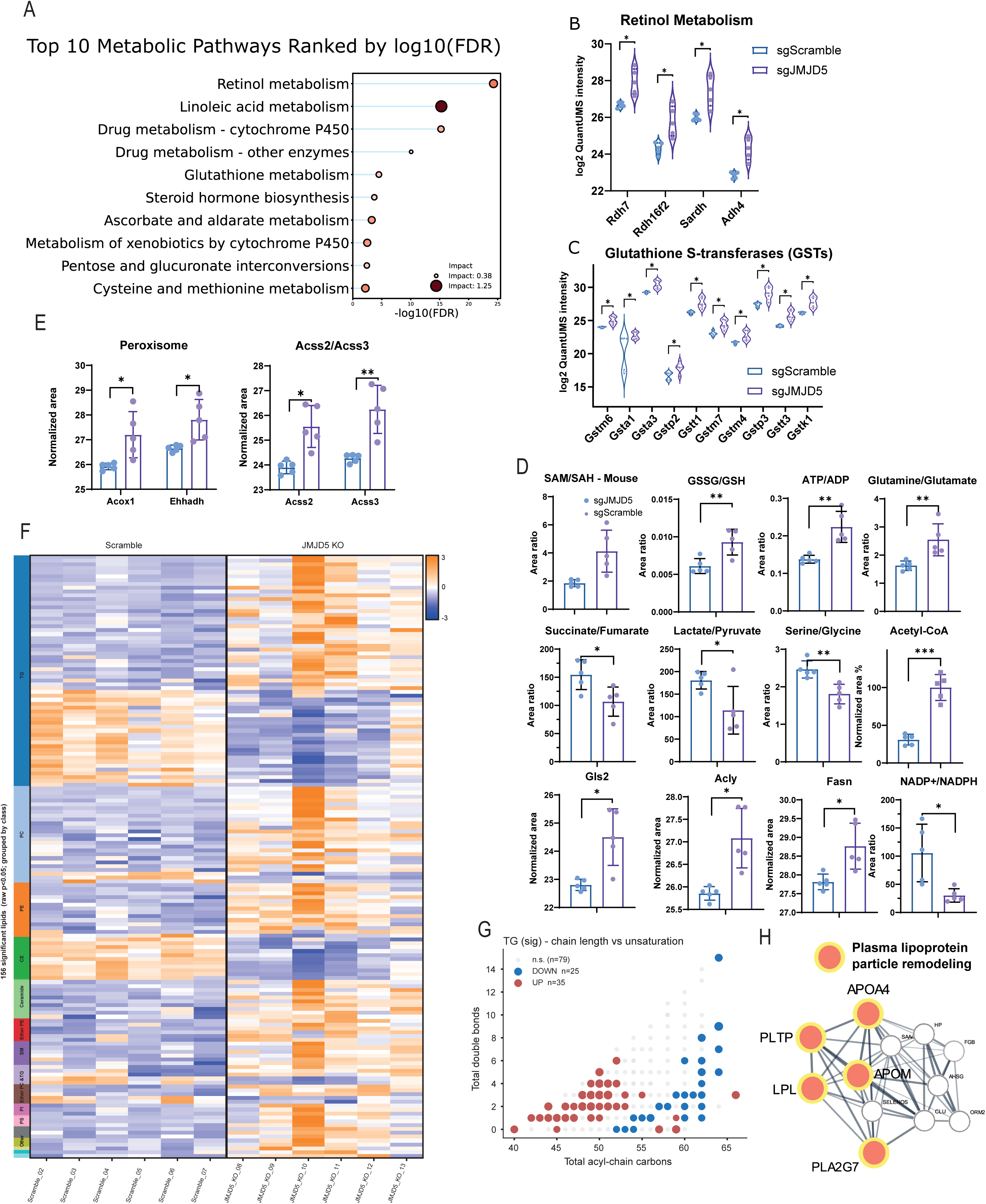
JMJD5 loss disrupts hepatic metabolism in vivo. Mouse liver tumours from sgScramble versus sgJMJD5 animals were profiled by proteomics (n = 6 and 5), metabolomics and lipidomics (n = 6 per group). (A) Joint pathway analysis integrating significantly up-regulated proteins (p < 0.05) with significant metabolite changes (p < 0.05), showing the top 10 metabolic pathways ranked by −log10(FDR). (B) Expression levels of enzymes of retinol metabolism (violin plots, sgScramble vs sgJMJD5). (C) Expression levels of glutathione S-transferases (GSTs) (violin plots, sgScramble vs sgJMJD5). (D) Grid of metabolite ratios and metabolic-enzyme levels in sgScramble vs sgJMJD5 livers (mean ± s.d.; significance by t-test) (E) Peroxisomal β-oxidation and acetyl-CoA-synthesising enzymes increased on JMJD5 loss: normalised levels of Acox1 and Ehhadh (first and second steps of peroxisomal VLCFA β-oxidation) and of Acss2 and Acss3 (cytosolic and mitochondrial acetyl-CoA synthetases) (mean ± s.d.; p < 0.05, p < 0.01). (F) Class-grouped heatmap of significantly changed mouse-liver lipid species (sgScramble vs sgJMJD5; raw p < 0.05; z-normalised), summarising the lipidome remodelling. (G) Triacylglycerol (TG) chain-length analysis: significant TG species plotted as total acyl-chain carbons (x) against total double bonds (y), coloured by direction of change. (H) STRING cluster of down-regulated secreted proteins enriched for the GO term “plasma lipoprotein particle remodelling” (Apoa4, Pltp, Apom, Lpl, Pla2g7), indicating coordinated suppression of lipoprotein metabolism.

Quantification of metabolite ratios revealed significant increases in SAM/SAH, GSSG/GSH, ATP/ADP, PCr/Cr, and glutamine/glutamate in the JMJD5-deficient group (Fig. 7 D). These changes reflect elevated methylation potential, increased oxidative stress, and heightened energy potential. Additionally, proteomic changes revealed the upregulation of Gpx1, consistent with increased peroxide detoxification. The increase in glutamine/glutamate, alongside Gls2 upregulation (Fig. 7 D), suggests elevated glutamine uptake and/or accelerated downstream glutamate utilisation for TCA cycle anaplerosis or glutathione synthesis. Conversely, the NADP⁺/NADPH, succinate/fumarate, and serine/glycine ratios were reduced (Fig. 7 D), consistent with altered TCA cycle dynamics, and altered one-carbon flux.

Proteomic analysis revealed upregulation of urea cycle enzymes Arg1, Ass1 and Asl (Data Table 8), and the metabolic profiling confirmed perturbations in amino acid metabolism, with L-arginine and L-cystine exhibiting the most severe depletion. Arginine reduction aligns with increased flux through the urea cycle, while cystine reduction may reflect redox stress and increased glutathione demand. The serine/glycine axis was also disrupted, with reductions in both serine and glycine, suggesting a broader inhibition or rerouting of one-carbon metabolism. The urea cycle intermediate citrulline was slightly depleted, while guanidinosuccinic acid, a marker of elevated nitrogen flux, was strongly elevated, reinforcing the activation of nitrogen disposal mechanisms.

Together, the significant increase in acetyl-CoA levels (Fig. 7 D) and the upregulation of ATP citrate lyase (Acly) (Fig. 7 D) indicate enhanced cytosolic conversion of citrate to acetyl-CoA, substrate for histone acetylation and fatty acid synthesis, the latter supported by the upregulation of fatty acid synthase (Fasn) (Fig. 7 D), fuelling lipid remodelling.

Within the TCA cycle, intermediates including malate, succinate, and fumarate were diminished. These data suggest a metabolic rewiring where glutaminolysis and amino acid catabolism are activated, yet TCA cycle intermediates were reduced despite the lower succinate/fumarate ratio and high ATP/ADP, suggesting a functional ETC and cataplerosis.

### JMJD5 deficiency remodels the lipidome

Lipidomic profiling uncovered a systematic shift in lipid composition on JMJD5 deletion. Amongst phospholipids, phosphatidyl choline (PC), phosphatidylethanolamine (PE) and phosphatidyl inositol (PI) species were broadly increased (Fig. 7 F, Extended Data Fig 7 A). However, the very long-chain polyunsaturated PC 44:6, PC 44:10, and PE 40:5 were selectively decreased (Extended Data Fig. 7 B). Phosphatidylethanolamine N-methyltransferase (Pemt), which uses SAM to methylate PE to form PC was significantly elevated, linking one-carbon metabolism to phospholipid biosynthesis. Cardiolipins (CL), especially with high linoleic acid content CL(18:2), were elevated, consistent with mitochondrial membrane adaptation to oxidative stress (Extended Data Fig. 7 C). Sphingomyelins (SM) and ceramides (Cer) were also increased (Extended Data Fig. 7 D & E). Cholesteryl esters (CE) were significantly reduced (Extended Data Fig. 7 F), and triacylglycerols (TG) exhibited a chain-length–dependent pattern with an increase in shorter-chain TGs, while long-chain TGs were consistently decreased (Fig. 7 G, Extended Data Fig. 7 G). The decrease in very long chain fatty acids (VLCFA) led us to interrogate the proteomics data, and we found significantly increased levels of Acox1 and Ehhadh (Fig. 7 E), which catalyse the first and second step in the peroxisomal oxidation of VLCFA, respectively. The peroxisomal β-oxidation of VLCFA exports acetate ^41^, later converted to acetyl-CoA in the cytosol by Acss2, and in mitochondria by Acss3 ^42^, both found to be increased (Fig. 7 E), revealing a broad effort to deal with peroxisomal and mitochondrial products. Furthermore, network cluster analysis using the down-regulated proteins revealed a group of apolipoproteins and lipid-metabolising enzymes (Apoa4, Pltp, Apom, Lpl, Pla2g7) enriched for the GO biological process “plasma lipoprotein particle remodelling” (Fig. 7 H), indicating coordinated suppression of lipoprotein metabolism in JMJD5-deficient liver tumour. Together, the lipidomics data revealed a JMJD5-dependent program regulating phospholipid remodelling, VLCFA peroxisomal β-oxidation and fatty acids de novo biosynthesis.

## Discussion

Mutations in 2-oxoglutarate-dependent dioxygenases (2OGDDs) have been linked to various genetic diseases. For example, mutations in prolyl-hydroxylases are directly linked to diseases through the misregulation of collagen hydroxylation ^43–45^, providing a clear connection between the loss of hydroxylase activity and a resulting phenotype. Mutations in JMJD5 cause severe phenotypes ^8^, but the direct mechanistic link between the genotype and phenotype has remained elusive. Our results reveal a novel signalling pathway that helps explain how inhibiting JMJD5 impacts the proteome and metabolome.

Our data suggest that many of JMJD5’s effects are transduced via a signalling pathway that requires its hydroxylase activity. Specifically, we’ve shown that hydroxylation-dependent complex formation between ISY1 and PRMT6 is a key step. When hydroxylation is reduced, ISY1 fails to sequester PRMT6, which in turn increases PRMT6 activity. This is likely the pathway’s “branch point,” as numerous proteins are regulated by asymmetric dimethylarginine (ADMA) modifications that are introduced by PRMT6 ^46^. Intriguingly, there’s a significant overlap between the phenotypes of JMJD5 loss and PRMT6 activation, suggesting a causal link between these proteins in a broader physiological context. The finding that PRMT6 is a downstream effector of JMJD5 loss, suggests that tumours with low JMJD5 levels may be susceptible to PRMT6 inhibitors, which have been developed as anti-neoplastic agents ^47^. This is something we will investigate in future work, along with the precise mechanism by which the interaction of ISY1 regulates PRMT6, including whether hydroxylated ISY1 affects PRMT6 activity in vitro, and how ADMA post-translational modifications transduce the effect of JMJD5 loss.

There have been controversies regarding whether some JmjC 2OGDD act on histones. The finding that JMJD5 regulates histone methylation status via regulating PRMT6 activity raises the question as to whether related indirect mechanisms may, at least in part, explain the apparent cellular links between the activities of other JmjC 2OGDD, including human JMJD4, 6, and 7 as well as the ribosomal protein hydroxylases MINA53/NO66, to histone modifications ^48^.

The strong phenotype observed from the loss of JMJD5 in our genetically engineered mouse model (GEMM) was unexpected. It dramatically curtailed liver tumour growth, which contradicts what has been observed in human liver tumours, where JMJD5 is frequently reduced and low expression is correlated with poor survival ^14^. One possible explanation is the very rapid nature of phenotypic effects in our model, which required animals to be culled just two weeks after hydrodynamic tail vein injection. As loss of JMJD5 has been shown to induce replication stress and reduce DNA-damage repair ^8^, this may explain why a rapid and highly proliferative tumour model is inhibited by JMJD5 loss. Alternatively, JMJD5 levels might have a “Goldilocks” value that is conducive to tumour growth, and a complete knockout could be anti-tumorigenic. We also note a paradox: the ATP/ADP ratio is higher in JMJD5-KO tumours despite their lower tumour burden (Fig. 7 D); we report this observation as a topic for follow-up studies.

2OGDDs are a broad enzyme superfamily whose reliance on 2OG, iron, and molecular oxygen appears to make them especially well-suited to sense the cell and organism’s metabolic and oxygenation states. In this study, we investigated JMJD5’s function using knockout experiments, which provided a broad overview of the signalling pathways it can regulate. While near-complete loss of JMJD5 and hence its hydroxylation activity has been shown to occur in genetic diseases as well as hepatic and pancreatic tumours, the precise physiological roles of JMJD5 catalysed hydroxylation remain to be addressed.

A notable feature of the 2OGDD superfamily is that some branches, such as the PHD/EglN prolyl-hydroxylases, act as exquisite oxygen availability / hypoxia sensors, in a manner proposed to reflect their slow reaction with, and *K_m_* values for, molecular oxygen ^49,50^, the latter of which fall within the physiological normoxic to hypoxic range ^2^. Other 2OGDDs are differently sensitive to changes in metabolite levels, including inhibition by their product succinate, fumarate or substrate 2-oxoglutarate (2OG). Peptide-based assays with JmjC 2OGDD are often conducted with 20 to 25-mer peptides, with the potential acceptor amino acid typically in the central region of the peptide. When we assayed the hydroxylation efficiency of isolated JMJD5, we noticed that a relatively short 15 residue ISY1 peptide fragment was a poor substrate compared to a longer 30 residue substrate, suggesting that more distant or secondary structure interactions play a role in substrate recognition by JMJD5, as is the case for at least some other JmjC 2OGDDs, including JMJD4 ^51^. Although we don’t have detailed kinetic parameters for JMJD5 with ISY1 protein substrates, we do have data for the related JmjC hydroxylase FIH. Although, like the PHD/EglN prolyl-hydroxylases, FIH is a HIF hydroxylase, its activity is much less sensitive to reduced oxygen levels, both in isolated form and in cells. FIH activity, like some other JmjC 2OGDD, is sensitive to inhibition by the (R)- and (S)- isoforms of 2-hydroxyglutarate (2HG) ^2^. *R*-2HG is an oncometabolite produced at low levels under normal conditions but is strongly induced by oncogenic mutations of isocitrate dehydrogenase (IDH1 and IDH2) ^52^. *S*-2HG, on the other hand, can be generated by malate dehydrogenase or under hypoxic and acidic conditions by lactate dehydrogenase, which catalyses the reduction of 2OG to S-2HG ^53^.

JMJD5 is known to be inhibited by both (S)- and (R)-2HG ^54^. The induction of *S*-2HG, driven by lower pH and high LDH activity due to prolonged hypoxia, could be what drives the metabolic adaptations we observed upon JMJD5 loss. While not investigated in detail here, we observed that inactivating JMJD5 induces proteomic changes that indicate it may drive cellular metabolism toward glucose consumption and, rather than shunting carbons toward lactate secretion, promote lipogenesis. This is a process that can generate ATP without oxygen consumption, similar to glycolysis. These observations are related to those with genetic and small molecule interventions of FIH, where effects on glycolysis, oxidative metabolism and lipid metabolism were all observed ^55^.

In the liver, our study reveals that JMJD5 loss affects retinol metabolism and peroxisomal β-oxidation, extending beyond traditional one-carbon metabolism and methylation dynamics. This shift is marked by increased retinal biosynthesis and retinoic acid (RA) production, which activates nuclear receptors involved in fatty acid oxidation. Enhanced PPARα signalling, evidenced by upregulated enzymes like Acox1, reflects increased peroxisomal activity. Lipidomic analysis shows a reduction in specific phospholipids, indicating more peroxisomal chain shortening and altered fatty acid flux. This is accompanied by heightened acetyl-CoA and Fasn levels, suggesting redirected carbons from oxidation into lipogenesis and membrane biogenesis. 2OGDD are known to play established roles in lipid metabolism via carnitine biosynthesis and α-oxidation of lipids, including phytanic acid, in the peroxisome ^56^. The more recent links between FIH and, as reported here, JMJD5 to lipid metabolism further extend the roles of 2OGDD in this regard ^55^. Furthermore, JMJD5-deficient tumours exhibit increased glutathione S-transferases (GSTs) linked to oxidative defence, potentially through retinoid–PPAR signalling. Network analysis reveals reduced HDL biogenesis, shifting lipid handling intracellularly and promoting remodelling.

Overall, JMJD5 loss initiates a retinoid-driven metabolic state enabling oxidative stress management and fatty acid turnover, impacting on membrane integrity and organelle function. This raises the intriguing possibility that JMJD5, through PRMT6 inhibition, may restrain retinoid metabolism and peroxisomal β-oxidation, maintaining a balance between lipid catabolism, methylation potential, and redox homeostasis. Loss of this restraint could predispose cells to altered lipid composition and turnover, with potential consequences for the health of the organ and predisposing it to metabolic disfunction. Whether this metabolic readjustment is the driving factor of reducing JMJD5 during liver progression is a topic that will have to be investigated further.

## Methods

### Reagents

All reagents were from Sigma unless otherwise stated. Solvents were LC-MS grade

### Cell lines

HEK293t were grown in DMEM 4.5 g/l glucose supplemented with 10% foetal bovine serum and 2mM glutamine, at 37°C and 5% CO2. HepG2 were grown similarly, but using 1g/l glucose

### Plasmids

Flag-myc-ISY1 was purchased from Origene. JMJD5 plasmids (wt and H321A) were a kind gift from Yoshihiro Izumiya ^13^. Flag-RCCD1 GFP-KDM* was a kind gift from Profs Sun and Wang ^57^,

### CRISPR/Cas9

**Table 1:**
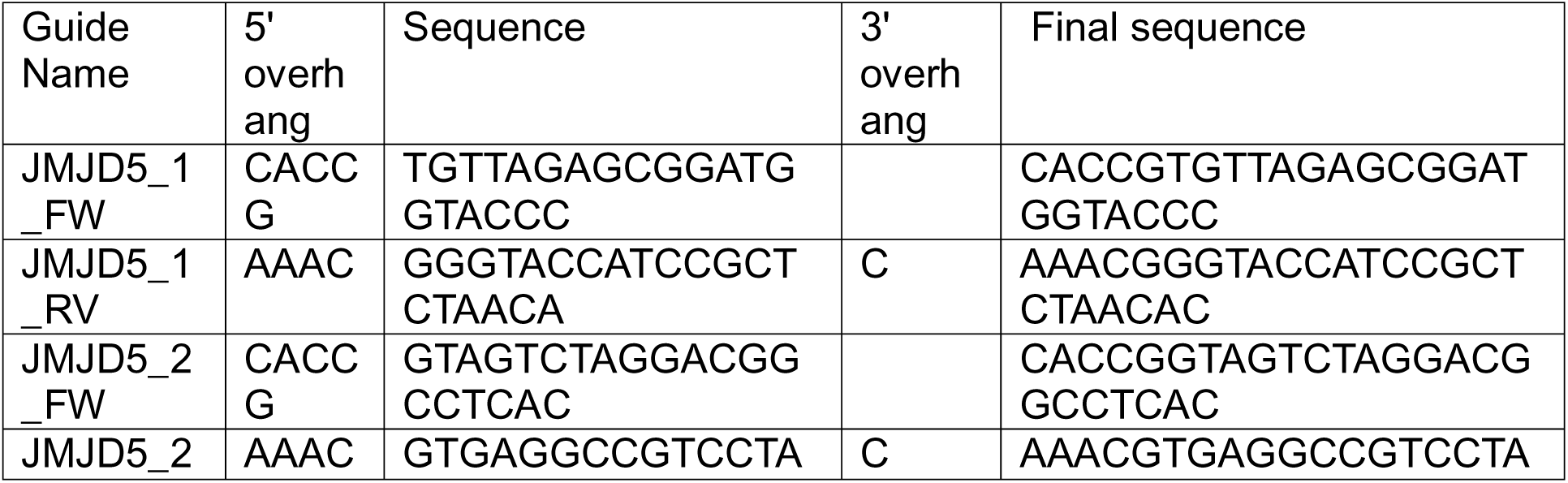
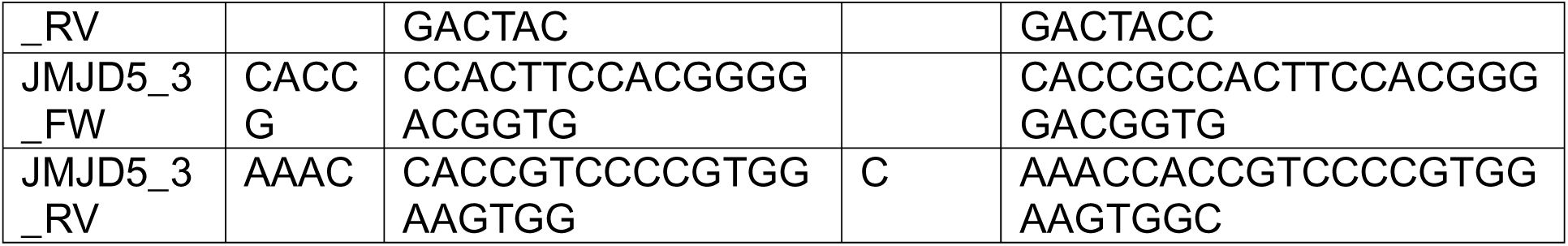
gRNA guides targeting mouse Jmjd5.

### Primary antibodies

| Serial No. | Catalogue No. | Name | Target Protein | Species reactivity | Host | Dilution | Dilution Medium (5% ) | Supplier |
| --- | --- | --- | --- | --- | --- | --- | --- | --- |
| 1 | 13522 | Asymmetric Di-Methyl Arginine Motif [adme-R] MultiMab® | meARG | All | Rabbit | 1:1000 | BSA | CST |
| 2 | Ab121250 | Anti-ISY1 | ISY1 | Human, mouse, rat | Rabbit | 1:100 | RIPA | Abcam |

### Western blot analysis

Cleared lysates were resolved on SDS-PAGE acrylamide gels and transferred onto PVDF membranes (Whatman) using Mini Trans-Blot® Electrophoretic Transfer Cell (BIO RAD). Membranes were blocked in 4% BSA for 1h and blotted with primary antibodies (see paragraph below). Immunocomplexes were visualized using ClarityTM Western ECL Substrate (BioRAD) in a ChemidocTM MP (BioRAD) with horseradish peroxidase–conjugated secondary anti-bodies (CST 1:10000). WB quantification was performed using ImageJ. All immunoblots experiments were repeated at least three times (n=3).

### Gene editing in cells

crRNAs (Alt-R CRISPR-Cas9 crRNA) targeting the genes of interest were designed and ordered from IDT (Table #) along with tracrRNA (Alt-R CRISPR-Cas9 tracrRNA) Cas9 (Alt-R™ S.p. Cas9 Nuclease V3; cat# 1081058) and electroporation enhancer (Alt-R™ Cas9 Electroporation Enhancer, 2 nmol; cat# 1075915). Equimolar concentrations of the tracrRNA and crRNA were mixed in a sterile PCR tube and duplexes were formed by heating at 95°C for five minutes then allowing the mixture to cool to room temperature. The ribonucleoprotein complex (RNP) was formed by adding 100pmol Cas-9 and 100pmol electroporation enhancer to 120pmol of the duplex and incubating it for ten minutes at room temperature.

Cells were grown to 80% confluency in a suitable dish, trypsinised, washed with PBS and counted. Nucleofection was carried out using the SF Cell Line 4D-Nucleofector® X Kit (V4XC-2032). 0.1x106 cells were suspended in 20uL SF Cell Line Nucleofector® Solution supplemented with Supplement 1 and pipetted into the Nucleocuvette® Strip. The RNP was pipetted into the cell suspension and the mixed carefully.

The RNP was transfected into the cells using the Lonza 4-D Nucleofector® (AAF-1003X) using the EH-100 program. The cells were immediately supplemented with 80uL fresh media and pipetted into 6-well plates. Cells were incubated for 48-72 hours with the media being changed after 24 hours. Knockout was confirmed by western blot, whole expression proteomics or IHC.

### CRISPR guides table

**Table 2:**
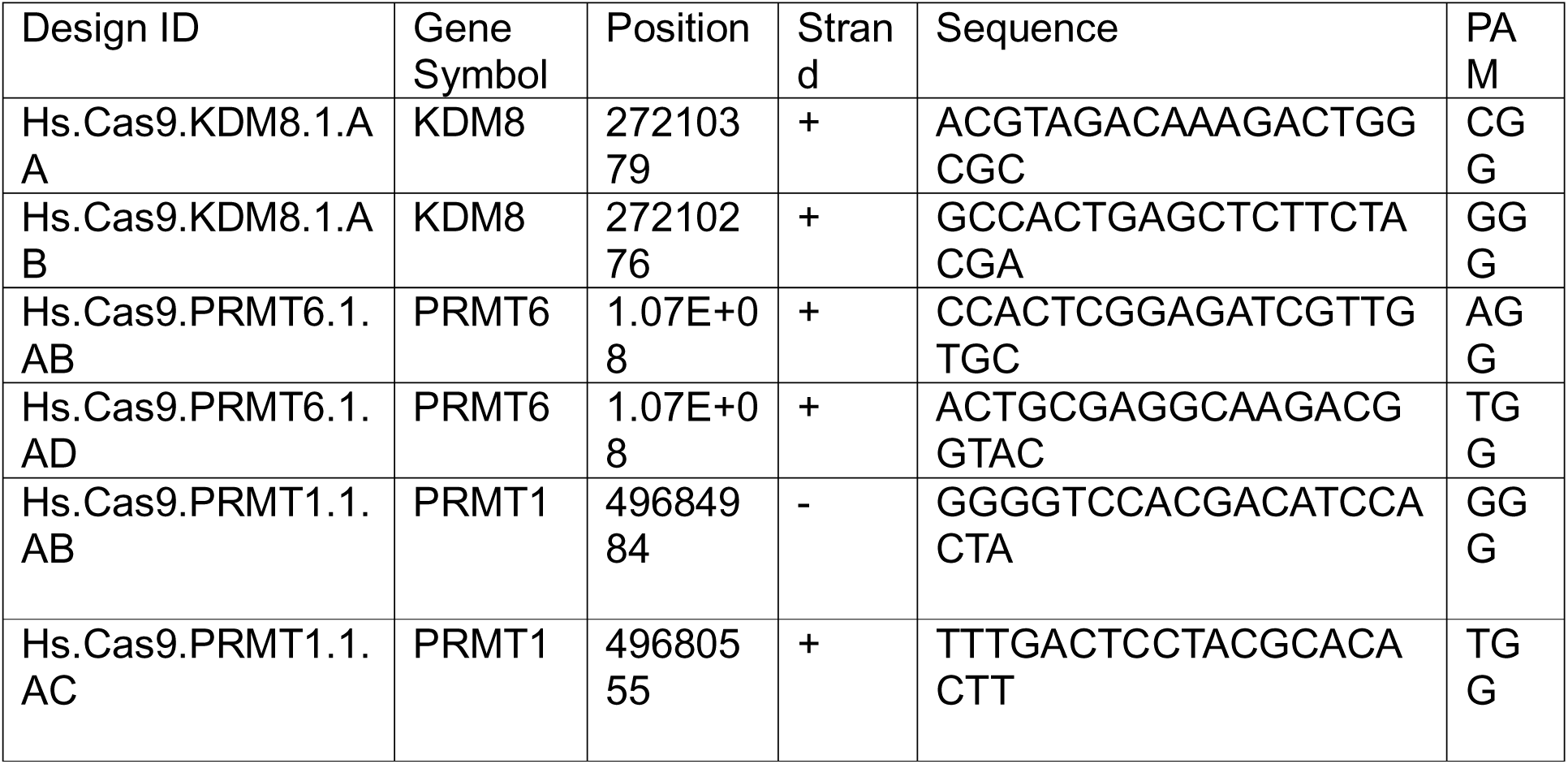
gRNA guides targeting human *Jmjd5*, *prmt6*, and *prmt1*.

### Animal experiments

Short guides (sg) sequences targeting the Trp53 gene (p53-SBCrispr), Jmjd5 gene (JMJD5-SBCrispr) (3 unique guides per gene, see table 1) or non-targeting short guides (Scramble-SBCrispr) were cloned into CRISPR-SB plasmid (Addgene #177936). Female FVB/N mice from Charles River (4-6 weeks of age) were given hydrodynamic tail vein injections containing DNA plasmids in physiological saline solution at a volume of 10% bodyweight (maximum 2 ml) as previously published ^58^. Mice were divided into two groups of 6 mice each. They were injected with solutions containing pCMV(CAT)T7-SB100 (6 µg, Addgene #34879), pT3-N90-beta-catenin (20 µg, Addgene #31785), p53-SBCrispr (20 µg), c-myc-PT3EF1a (20 µg, Addgene #92046) and Scramble-SBCrispr (20 µg) (sgScramble) or JMJD5-SBCrispr (20 µg). After 2 weeks, mice were euthanised by CO2 and death was confirmed by dislocation of the neck. Livers were perfused with PBS. The liver lobes were separated, and the left lobe was fixed with 4% paraformaldehyde overnight. The right, median and caudate lobes were dissected and frozen using isopentane cooled to -78.5°C using dry ice.

### Immunohistochemistry

Formalin-fixed, paraffin-embedded mouse tumour sections were stained for p53, MYC, β-catenin (CTNNB1), γ-H2AX and ADMA. Sections underwent heat-mediated antigen retrieval, were incubated with the primary antibody and detected with a biotinylated secondary antibody; antigen-retrieval conditions and antibody dilutions are detailed in the antibody table below.

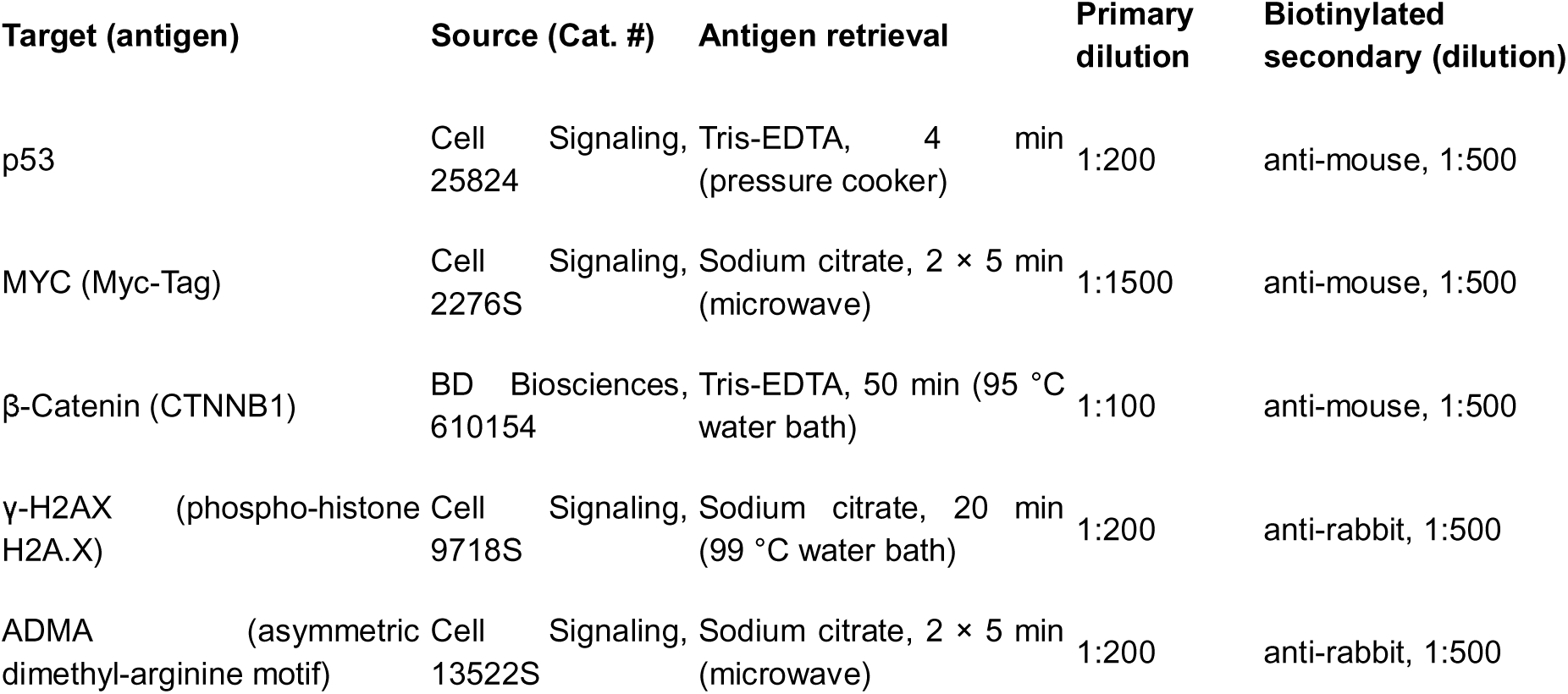

### Generation of stable fluorescent reporter cell lines

To reconstitute JMJD5/PRMT6 double-knockout (dKO) HepG2 cells JMJD5 and PRMT6 were re-introduced by viral transduction. eGFP-JMJD5 was delivered with a standard (Moloney-based) retrovirus, and V5-mKate2-PRMT6 with a second-generation lentivirus Both viruses were produced in Lenti-X HEK293T cells. dKO cells were first transduced with the eGFP-JMJD5 retrovirus (for double-reconstituted lines) or left untransduced, then transduced with the V5-mKate2-PRMT6 lentivirus by spinfection in the presence of polybrene and selected with puromycin. This yielded “JP” cells re-expressing both JMJD5 and PRMT6 and “P” cells expressing PRMT6 only.

### Super-resolution microscopy

eGFP-JMJD5 together with V5-mKate2-PRMT6 (“JP” cells) or V5-mKate2-PRMT6 alone (“P” cells were seeded onto rat-tail-collagen-coated high precision cover-glasses (Marienfeld, Germany). Cells were fixed in 4% paraformaldehyde for 10 min, washed, quenched and permeabilised with 0.3% Triton X-100 for 5 min. Endogenous ISY1 was detected by overnight incubation with an anti-ISY1 antibody (1:100; Bethyl Laboratories / Thermo Fisher Scientific) in TBST. The following day, the coverslips were washed and incubated with ChromoTek GFP-Booster (ChromoTek) and an anti-rabbit Alexa Fluor 647 (Thermo Fisher) secondary antibody. mKate2-PRMT6 was imaged by its intrinsic fluorescence. Coverslips were mounted in an aqueous, DAPI-containing mountant (Thermo Fisher Scientific) and imaged by super-resolution microscopy. Co-localisation between ISY1 and PRMT6 was quantified as a co-localisation coefficient across multiple fields. Images were acquired using Nikon SoRa ™ system (Nikon Europe B.V.). Imaging was carried out using a SR Apo TIRF 100x 1.49NA oil lens (Nikon Instruments). The CMOS cameras used for acquisition were Teledyne Photometrics Prime 95B (Teledyne Photometrics 3440 E.Britannia Drive, Tucson AZ) and 405 / 488 / 561 / 640nm laser lines. Z-step size for Z stacks was set to 0.100 um as required by manufacturers software. Dapi was captured using a 405 laser and 447/60 filter, JMJD5 a 488 laser and a 525/50 filter, ISY1 a 640 laser and a 708/75 filter, and PRMT6 a 561 laser and a 600/52 filter. Acquisition of images, image registration and deconvolution was carried out using the 3D automatic algorithm with Nikon NIS Elements Advanced Research software (ver 6.10.01). Figures were prepared using FIJI (1.54p) and colocalisation was measured by using JACoP plugin ^59^

### Expression Proteomics

HepG2 cells and liver protein pellets were processed following the PAC digest protocol ^60^. Cells and protein pellets were lysed with PAC lysis buffer (5% SDS, 100 mM Tris pH 8.5, 1 mg/ml chloroacetamide, 1.5 mg/ml Tris (2-carboxyethyl) phosphine). They were sonicated and then heated at 95 degrees for 30 minutes.

The lysates were then added to a Kingfisher 96 well deep well plate (Thermo Fisher, UK) and prepared for an 8h digest protocol on the Kingfisher Duo (Thermo Fisher, UK). Lysate was added to Row G with HILIC beads (MagReSyn, UK) and MeCN was added to 70%v/v final concentration. Rows D, E, and F were filled with 95% v/v MeCN and Rows B and C were filled with 70% v/v EtOH. The digest buffer (1 ug/ml MS grade trypsin in 50 mM triethyammonium bicarbonate) was added to Row A.

Desalted peptides were then loaded onto 25cm Aurora Columns (IonOptiks, Australia) using a RSLC nano uHPLC systems connected to a Fusion Lumos mass spectrometer. Peptides were separated by a 70 min linear gradient from 5% to 30% acetonitrile, 0.5% acetic acid. The mass spectrometer was operated in DIA mode, acquiring a MS 350-1650 Da at 120k resolution followed by MS/MS on 45 windows with 0.5 Da overlap (200-2000 Da) at 30 k with a NCE setting of 28. Files were searched using DIA-NN 1.8 or 2.0 ^61^ against the human Uniprot database and a list of common contaminants using pre-set settings. Statistical test and analysis was done on Perseus ^62^.

### Interaction Proteomics

Cells were lysed in lysis buffer (1% NP40, 150 mM NaCl, 2 mM EDTA, 1 mM PMSF, 20 mM Tris pH 7.5). The lysates were cleared by centrifugation and pulldowns were performed in a Kingfisher Duo (Thermo Scientific). Cleared lysates were incubated with 5µL of protein G Mag Sepharose™ Xtra beads (GE Healthcare) and 1µg of antibody anti-ISY1 or antibodies against EGFP, Myc (both Chromotek, UK) or Flag (Sigma, UK)pre-coupled to magnetic agarose beads for 1 hour, washed two times with Lysis Buffer and three times with TBS then resuspended in 100 µl digestion buffer (2 M urea, 50 mM Tris-HCL pH7.5, 1 mM DTT) with 0.25µg of porcine trypsin MS-grade (Promega). Samples were digested at 37°C for 8 hours. Cysteines were alkylated with iodacetamide (1mg/ml), 30min. and acidified with TFA (final concentration 1% v/v). Peptides were desalted and purified using C18 STAGE tips. The eluted peptides were lyophilised in a vacuum concentrator (LabConco), resuspended in 0.1% TFA and analysed by LC-MS/MS on a Fusion Lumos Mass Spectrometer (Thermo Fisher) using OT/OT DDA acquisition, 120k MS and 30k MS/MS resolution.. Label-free quantification approach was used for protein identification and quantification using FragPipe software ^63^.

Alternatively, for the ISY1 interactome, LC-MS/MS analyses were performed on a timsTOF SCP Mass Spectrometer (Bruker) coupled to an Evosep One LC system (Evosep). We utilised Aurora Elite CSI analytical columns (IonOpticks, AUR3-15075C18-CSI) and a CaptiveSpray ionisation source. Samples were analysed using the Whisper 40 SPD zoom method, which employed a gradient flow of 200 nL/min with a 31 min method duration. Eluted peptides were analysed with a parallel accumulation-serial fragmentation data-independent acquisition (diaPASEF) method. Isolation windows were optimised for low-input samples using an open-source Python package for dia-PASEF methods with Automated Isolation Design (py_diAID) ^64^ Data acquisition: employed a window placement scheme consisting of 2 TIMS ramps with 12 mass ranges per ramp spanning from 327 to 1200 m/z and from 0.6 to 1.60 1/K0 and searched using Spectronaut 20 (Biognosis).

### Crosslinking Mass Spectrometry

Immunoprecipitated complexes were incubated with PhoX (Thermo Fisher) at 2 μg/ml in PBS and rotated at RT for 1 hour. Crosslinking was quenched using 50 mM ammonium bicarbonate for 5 minutes and digested as described above. Crosslinked peptides were enriched as described ^24^ but using Zr IMAC beads (MagResyn) rather than Fe beads. Enriched peptides were loaded onto Evotips and analysed on a timsTOF SCP as described above, but using a DDA PASEF method with 200 ms accumulation, 10 PASEF elutions and a window excluding likely singly charged peptides. The raw data were searched against the human proteome in Fragpipe ^25^ to generate a calibrated mxXML that was converted into mgf using ProteoWizard MSConvert ^65^. The spectra were then analysed using xiSEARCH ^66^ with MS1/MS2 error tolerances of 14 and 15 ppm, respectively. The search was performed with carbamidomethylation of cysteine as a fixed modification and methionine oxidation as a variable modification. The PhoX cross-linker was defined to be reactive with K,S,T,Y residues and protein N-termini, with a score penalty for matches to S,T,Y residues. Modifications related to the cross-linker included linked PhoX (+209.97181 Da), hydrolysed PhoX (+227.98237 Da) and amidated PhoX (+226.99836 Da), searched on protein N-termini and K,S,T,Y residues of linear peptides. Results were then filtered in xiFDR to a 5% false discovery rate at the residue pair level and exported to xiView.org for visualisation.

### Histone PTM analysis

Histones were purified from HepG2 cells as described ^36^. Histones were extracted by acid extraction using 0.2 M sulfuric acid. The extracted histones were propionylated and digested with trypsin using a modified bottom-up proteomic approach. Briefly, the histones were first reacted with propionic anhydride to block lysine N□-amino groups. Following an overnight trypsin digestion, the resulting peptides were subjected to a second propionylation step to modify the newly formed N-termini. The peptides were then purified and analysed by LC-MS/MS on a Fusion Lumos Mass Spectrometer (Thermo Fisher) using OT/OT DIA acquisition, 60k MS and 30k MS/MS resolution and 50 Da windows. Data was analysed using the Matlab Epiprofile script.

### Extraction of protein, metabolites and lipids from the liver

Fresh frozen liver tissue was weighed and homogenised in methanol (1:20 w/v) using the soft tissue setting on the Precellys homogeniser (cat# P002511-PEVT0-A.0). Methyl tert-butyl ether (MTBE) was added to the methanol in a 3:1 ratio and centrifuged at maximum speed at 4°C for 10 minutes. The supernatant was moved to a new tube, and the pellet was lysed using lysis buffer (5% SDS, 100 mM Tris-HCl pH 8.5, 1 mg/ml CAA, 1.5 mg/ml TCEP), proteins were extracted using the PAC digest protocol mentioned above and LC-MS/MS was performed as mentioned above.

LC-MS grade water was added to the supernatant (1:1 water to methanol) and vortexed to mix. The tube was centrifuged at max speed at 4°C for 10 minutes. The solution, now separated into two phases, was pipetted put carefully to collect only the upper phase and moved to a new tube. The lower phase, now containing metabolites was chilled at -80°C for ten minutes to precipitate any remaining protein. The tube was then centrifuged at max speed at 4°C for 10 minutes. 30-50uL of supernatant was used for LC-MS as mentioned above.

The upper phase, now containing lipids, was dried using a vacuum centrifuge at room temperature. The dried lipids were reconstituted in minimum volume (100mL) of 1:1 methanol: butanol v/v. The tube was chilled at -80°C for ten minutes to precipitate any remaining protein. The tube was then centrifuged at max speed at 4°C for 10 minutes. 30-50uL of supernatant was used for LC-MS as mentioned above.

### LC-MS

ZIC-pHILIC LC column with Q Exactive or Q Exactive Plus MS (Short Run) were used. SeQuant ZIC-pHILIC guard column 20 x 2.1mm and SeQuant ZIC-pHILIC analytical column 150 x 2.1mm, 5µm were used. 20mM ammonium carbonate, adjusted to approximately pH 9.2 with ammonium hydroxide solution (25%) was used as mobile phase A and 100% acetonitrile for mobile phase B. Positive/negative mass spectrometry was used with no lock masses. The scan range was 70-9000 m/z, and the resolution was 70,000 with the maximum IT of 250ms.

### Data Analysis

The chromatogram was analysed, and the ratios were obtained using Skyline (MacCoss Lab Software). Metaboanalyst or Metabolite AutoPlotter was used to generate graphs.

#### Lipidomics

Lipids extracted from liver tissue (see “Extraction of protein, metabolites and lipids from the liver”) were analysed by LC-MS/MS and the resulting data used for the lipidomic profiling in Data Table 10. Extracts were analyzed by LC–MS using a Dionex LC system (Thermo Fisher Scientific) coupled to a Q Exactive Plus mass spectrometer (Thermo Fisher Scientific). Samples were acquired in both positive-and negative-ion modes in two separate runs. Chromatographic separation was performed on a Kinetex EVO column (150 × 1.0 mm, 1.7 µm; Phenomenex) maintained at 45 °C, using an autosampler held at 5 °C. Mobile phase A consisted of 60% acetonitrile and 40% methanol containing 5 mM ammonium formate and 0.1% formic acid, and mobile phase B consisted of isopropanol containing 5 mM ammonium formate and 0.1% formic acid. The flow rate was 0.075 mL/min and the total run time was 25 min. The gradient was as follows: 0–1.0 min, 0% B; 1.0–17.0 min, linear increase to 99% B; 17.0–21.0 min, 99% B; 21.0–21.5 min, return to 0% B; and 21.5–25.0 min, 0% B for re-equilibration. Lipidomics raw files (positive and negative mode, separate injections) were processed in Compound Discoverer 3.2 (Thermo Fisher Scientific) for peak detection, alignment, and integration. Lipid annotation was performed in LipidEx version 1.1.0, with in silico fragmentation matching against the LipidBlast, LipidEx HCD Formic, and LipidEx HCD Hydroxy spectral libraries. Positive- and negative-mode feature tables were merged into a single combined feature table prior to downstream analysis. The merged feature table was imported into MetaboAnalyst 6.0 and subjected to median normalisation and log2 transformation.

### RNA-sequencing and qPCR

Total RNA was extracted from HepG2 WT and JMJD5−/− cells using the QIAGEN RNeasy kit according to the manufacturer’s instructions. Approximately 1.5 × 10□ cells were seeded in 10 cm tissue culture dishes and grown to approximately 80% confluence. Cells were harvested to obtain a cell pellet, lysed, and processed according to the kit protocol. An on-column DNase digestion step was performed to remove residual genomic DNA. Purified RNA was eluted in RNase-free water, and RNA concentration and purity were initially assessed using a NanoDrop spectrophotometer. RNA-seq libraries were prepared from 500 ng total RNA per sample using the NEBNEXT Ultra II Directional RNA Library Prep Kit and the NEBNext Poly-A mRNA Magnetic Isolation Module on the NGeniuS automated platform, according to the manufacturer’s protocol. Sequencing was conducted on an Illumina NextSeq 2000 platform using a NextSeq 1000/2000 P2 XLEAP-SBS 200-cycle reagent kit. A P2 flow cell was used to run the libraries and sequenced using a 2 × 100 bp paired-end configuration. PhiX Control v3 was spiked into the run at 1%. Raw basecall data were converted to FASTQ files through Illumina BaseSpace. FASTQ files were processed using the Nextflow based nf-core/rnaseq pipeline version 3.18.0, from the nf-core/rnaseq GitHub repository. The pipeline was run with Nextflow version 24.10.5, build 5935. Reads were quality controlled and transcript abundance was quantified using Salmon. Transcript level estimates were imported into R using tximport and summarised to gene level length scaled counts. Differential expression between JMJD5−/− and WT HepG2 cells was assessed using DESeq2, with genotype included as the experimental variable. Genes with Benjamini–Hochberg adjusted P values < 0.05 were considered differentially expressed.

### In-vitro hydroxylation assays

Hydroxylation assays were performed in 96-well polypropylene plates (Agilent) and monitored by solid phase extraction coupled to mass spectrometry (SPE-MS). Purified recombinant full length JMJD5 (0.5 mM) was incubated with a synthetic ISY1 N-terminal peptide (5 mM) – either ISY1(2–16) (Ac-ARNAEKAMTALARFR) or ISY1(2–31) (Ac-ARNAEKAMTALARFRQAQLEEGKVKERRPF) – in the presence of L-ascorbate (100 mM), ferrous ammonium sulphate (10 mM) and 2-oxoglutarate (50 mM) for 1 h at room temperature. Enzyme reactions were performed in 50 mM HEPES pH 7.5 buffer. After 1 hour the enzyme reaction was quenched by addition of formic acid to a final concentration of 1% (v/v) and the plate transferred to an Agilent RapidFire RF365 high-throughput sampling robot connected to an Agilent 6550 ifunnel accurate mass quadrupole time-of-flight (QTOF) mass spectrometer. Samples were aspirated under vacuum and loaded onto a C4 SPE cartridge and the cartridge washed with 0.1% (v/v) aqueous formic acid (5.5 s at a flow rate of 1.5 ml min-1) to remove non-volatile buffer salts. The peptide was then eluted from the C4 SPE with 85% (v/v) MeCN and 15% (v/v) LCMS grade water and containing 0.1% (v/v) formic acid onto the mass spectrometer (5.5 s at a flow rate of 1.25 ml min-1). The mass spectrometer was operated in the positive electrospray ionization (ESI) mode with a drying gas temperature (280 °C), drying gas flow rate (13 L min-1), nebulizer pressure (40 psig), sheath gas temperature (350 °C), sheath gas flow rate (12 L min-1), capillary voltage (4000 V), nozzle voltage (1000 V), fragmentor voltage (365 V). Data were analyzed using Agilent Masshunter qualitative analysis (B.07.00). For the enzyme time course with ISY1(2–31), the enzyme reaction was performed in a deep well 96-well polypropylene plate in a total reaction volume of 1.0 ml. The RapidFire high-throughput sampling robot was programmed to take one sample from the enzyme reaction every 1.0 min. Extracted ion chromatogram data were extracted and integrated using Agilent RapidFire Integrator (version 4.3.0) and data plotted in GraphPad prism (version 5.04).

### Patient-cohort transcriptomic and proteomic analysis

JMJD5/KDM8 and RCCD1 transcript levels were compared across healthy liver, tumour-adjacent normal liver and primary hepatocellular carcinoma using the UCSC Toil re-processed RNA-seq compendium, Liver samples were partitioned into GTEx healthy liver (n = 110), TCGA-LIHC tumour-adjacent solid-tissue normal (n = 50) and TCGA-LIHC primary tumour (n = 369), and group distributions were compared by two-sided Mann–Whitney U test. Protein-level validation used the CPTAC HBV-related HCC proteogenomic cohort ^37^, JMJD5 cancer-cell-line mRNA was obtained from the Human Protein Atlas,

### Structural modelling

Structural models of the JMJD5/RCCD1/ISY1 and ISY1/PRMT6 complexes were generated with AlphaFold-3 using the AlphaFold Server (https://alphafoldserver.com) with default settings, submitting the protein FASTA sequences together with a Mn(II) ion at the JMJD5 active site. Predicted aligned error (PAE) and model confidence scores were used to assess the predictions, and models were visualised and rendered in UCSF ChimeraX.

### Network and functional-enrichment analysis

Protein–protein interaction networks were built in Cytoscape using the STRING app: all significantly regulated protein entries from a given cluster were imported as a STRING network, and Gene Ontology Biological Process (GO:BP) term enrichment was computed with STRING’s built-in functional-enrichment analysis. Network nodes were coloured by the enriched GO:BP categories.

### TCGA analysis

Normalised STAR-counts of RNA-seq data from the TCGA-LIHC study was used to stratify the data into two groups based on JMJD5 expression using the mclust script ^67^. This resulted in 281 low JMJD5 and 93 high JMJD5 samples. 5 samples in the low group and 1 sample in the high group were excluded for having high variance in the data. Mutational data from the study was analysed using the maftools package ^68^. 200 FLAG genes which were mutated in both groups were also eliminated from the analysis. The final results were 37 genes frequently mutated in JMJD5 low group (figure). Beta catenin (CTNNB1) was the most frequently mutated gene when JMJD5 was lost in hepatocellular carcinoma patients.

### Data analysis Statistics

Bioinformatic analysis was done on Perseus. Proteomics data was log2 transformed, proteins with x<3 missing values in at least one group were excluded. The remaining missing values were imputed using a random distribution as implemented in Perseus, statistical tests were either ANOVA or Student’s t-test, and multiple testing was corrected using permutation as implemented in Perseus.

### Data availability

The mass spectrometry proteomics data have been deposited to the ProteomeXchange with the dataset identifier PXD067356

## Acknowledgements

ZK is supported by the Melville Trust – For the Care and Cure of Cancer PhD studentship. AvK acknowledges the MRC Equipment grant (MR/X01293X/1), BBSRC Alert (BB/X019160/1) and Wellcome Trust Multi-user Equipment Grant (grant ID: 208402/Z/17/Z). EJ is supported by a CRUK BTP grant: (PRCBTP-May24/100001) and a PSC Support research 668 grant (EJPROJ23). A Cancer Research UK Fellowship (C52499/A27948 and PRCBTP-May24/100001) and MRC project grant (MR/Z506199/1) funds LB. He is also co-supported by a CRUK Centre Grant (CTRQQR-2021\100006). We thank Profs Michael Clague and Richard Scheltema, University of Liverpool, for fruitful discussions and technical advice. CJS thanks the Wellcome Trust (106244/Z/14/Z) and CRUK (CRUK/A24759) for support.

## Extended Data

Extended Data Fig. 1: JMJD5 expression in liver cancer and the transcriptional/one-carbon context of its loss. (A) JMJD5 mRNA expression (nTPM) across cancer cell lines from the Human Protein Atlas, (B) Volcano plot of the HepG2 proteome (JMJD5 −/− vs WT), the two-group companion to the four-way heatmap in Fig. 1B; significant proteins are colour-coded by category (JMJD5-KO confirmation, DNA-damage repair/JMJD5–RCCD1 complex, RNA metabolism, sulphur-compound/methionine metabolism, lipid metabolism). (C) RNA-seq of methionine/one-carbon-cycle genes in HepG2 WT versus JMJD5 KO (per-gene TPM; SHMT1, BHMT, BHMT2, MAT2B, MTHFR, MTR, MTRR, CBS), showing that the one-carbon programme is reproduced at the transcript level. (D) Proteome-versus-transcriptome correlation: protein log2 fold-change (JMJD5 KO vs WT) plotted against the corresponding mRNA log2 fold-change, showing that most JMJD5-driven proteome changes are poorly mirrored at the transcript level (sulphur/one-carbon metabolism being the main concordant exception). (E) Schematic of the methionine/one-carbon cycle, indicating the betaine arm (BHMT/BHMT2; betaine → homocysteine → methionine), the folate arm, and the SAM/SAH methylation node affected by JMJD5 loss.

Extended Data Fig. 2: Cross-linking MS/MS evidence for the ISY1 N-terminus at the JMJD5 active site. Representative annotated MS/MS spectrum of the inter-protein cross-linked peptide that links the acetylated ISY1 N-terminal peptide (M·A·R·N·A·E·K, residues around the R3 site; shown in purple) to a JMJD5 tryptic peptide (shown in green), identified by cross-linking mass spectrometry (PhoX cross-linker; xiSEARCH). Matched b- and y-ions for both peptide chains are labelled; the x-axis is m/z and the y-axis is intensity (right axis, % of base peak).

Extended Data Fig. 3: Non-canonical ISY1 N-terminal peptides and in-vitro hydroxylation of ISY1 by JMJD5. (A) Volcano plot of ISY1 peptides detected in HEK293 cells co-expressing ISY1 with either wild-type JMJD5 (JMJD5wt) or catalytically dead JMJD5 H321A (JMJD5mut), quantified with FragPipe (n = 3). (B) Bar graph (semi-LysC digestion) of three ISY1 N-terminal peptides — AMTALARFRQAQLEEGK, MTALARFRQAQLEEGK and TALARFRQAQLEEGK — in HEK293 cells co-expressing JMJD5wt versus JMJD5mut (log2 LFQ intensity; mean ± s.d., n = 3; p < 0.05). (C) Time course of in-vitro hydroxylation of the synthetic ISY1 N-terminal peptide ISY1□□□□ (Ac-ARNAEKAMTALARFRQAQLEEGKVKERRPF) by purified recombinant JMJD5. Reaction conditions: JMJD5 (0.5 mM) with L-ascorbate (100 mM), ferrous ammonium sulphate (10 mM) and 2-oxoglutarate (50 mM), at room temperature. (D) Mass spectrum of the synthetic ISY1 N-terminal peptide (Ac-ARNAEKAMTALARFR, ISY1□□□□) after 1 h in-vitro incubation with recombinant JMJD5, showing the +16 Da hydroxylated product (m/z 1746.90 → 1762.90). Reaction conditions as above, with 5 mM synthetic ISY1□□□□ peptide. Abbreviations: JMJD5wt/JMJD5mut, wild-type / H321A catalytically dead JMJD5; LFQ, label-free quantification; Ac, N-terminal acetyl; 2-OG, 2-oxoglutarate; ISY1□□□□ / ISY1□□□□, synthetic ISY1 peptides spanning residues 2–16 and 2–31.

Extended Data Fig. 4: AlphaFold model of the ISY1–PRMT6 complex. (A) AlphaFold model of the dimeric ISY1/PRMT6 complex. ISY1 Arg3 (R3) and the residues forming hydrogen bonds at the interface are shown as sticks (ISY1, pink/red; PRMT6, blue/purple). (B) Model-confidence diagnostics for the prediction in (A): the predicted aligned error (PAE) matrix and per-residue confidence, indicating a confidently predicted ISY1–PRMT6 interface. (C) Close-up of ISY1 R3 in the AlphaFold model of the dimeric complex containing hydroxylated ISY1-R3 (ISY1-OH-R3/PRMT6); residues involved in hydrogen bonds are shown, illustrating how R3 hydroxylation is accommodated at the PRMT6 interface.

Extended Data Fig. 5: PRMT1 knock-out and JMJD5/PRMT1 double-knock-out proteomic series. Genetic test of the alternative Type-I arginine methyltransferase PRMT1, profiled by mass spectrometry across four HepG2 lines: WT, PRMT1 −/−, JMJD5 −/−, and the JMJD5/PRMT1 double knock-out (DKO). (A) JMJD5 and PRMT6 protein levels by mass spectrometry across the genotypes, confirming line identity. (B) Heatmap of proteins significantly changed across WT, PRMT1 −/−, JMJD5 −/− and JMJD5/PRMT1 DKO (z-normalised per protein; triplicates). (C) Per-cluster rescue profiles (spaghetti plots) showing the behaviour of each protein cluster across the four genotypes; PRMT1 loss does not return the JMJD5-driven clusters to WT. (D) Proteome-wide perturbation per genotype, expressed as the number of significantly changed proteins relative to JMJD5 −/− (set to 100%). (E) QuantUMS intensities of JMJD5, PRMT1, BHMT, BHMT2 and CBS across the four lines (mean ± s.d., n = 3); in the JMJD5/PRMT1 DKO the one-carbon enzymes are not normalised (e.g. BHMT rises further), in contrast to the PRMT6 DKO. (F) ADMA Western blots for the knock-out lines.

Extended Data Fig. 6: Patient-cohort validation of JMJD5/RCCD1 loss and mouse-tumour immunohistochemistry. (A) JMJD5/KDM8 mRNA across GTEx healthy liver (n = 110), TCGA-LIHC tumour-adjacent normal (n = 50) and TCGA-LIHC primary tumour (n = 369), from the UCSC Toil re-processed compendium (log2 RSEM-normalised counts; two-sided Mann–Whitney U). (B) RCCD1 mRNA across the same three groups; RCCD1 transcript is modestly higher in tumour than in adjacent-normal (p = 2.2 × 10□¹□). (C) JMJD5/KDM8 protein in the CPTAC HBV-related HCC cohort (Gao et al., 2019; PDC000198; TMT-11-plex), paired tumour vs peri-tumour normal per patient (n = 155 pairs; median Δlog2(T/N) = −0.35; reduced in 80% of patients; paired Wilcoxon p = 4.5 × 10□¹□). Lines connect paired samples; bars are group medians; the inset histogram shows the per-patient Δ distribution. (D) RCCD1 protein in the same CPTAC cohort, paired (n = 165 pairs; median Δlog2 = −0.25; reduced in 77%; paired Wilcoxon p = 4.2 × 10□¹□), confirming co-ordinate protein-level loss of JMJD5 and RCCD1 in patient tumours. (E) Mutation co-occurrence analysis: forest plot / table of genes differentially mutated in JMJD5-low (n = 281) versus JMJD5-high (n = 93) TCGA-LIHC tumours (maftools; odds ratio with 95% CI; 37 genes enriched in the JMJD5-low group; p < 0.01). CTNNB1 (β-catenin) is the most frequently mutated gene when JMJD5 is low. (F) Immunohistochemistry of mouse tumour sections (sgScramble vs sgJmjd5) for p53, CTNNB1 (β-catenin) and MYC with a magnified MYC inset. (G) Jmjd5 protein level by mass spectrometry (sgScramble vs sgJmjd5), used to confirm knock-down. (H) γ-H2AX immunohistochemistry (sgScramble vs sgJmjd5), reporting increased DNA damage on JMJD5 loss. (I) Per-animal Western blots for asymmetric dimethyl-arginine (ADMA) with β-actin loading control, across three biological replicates (sgScramble vs sgJmjd5).

Extended Data Fig. 7: JMJD5 deficiency remodels the mouse-liver lipidome. Mouse liver lipidomics, sgJMJD5 versus Scramble (n = 6 per group); significance at raw p < 0.05 unless stated. (A) Lipid-class compositional shift: mean log2 fold-change (sgJMJD5/Scramble) per lipid class with s.e.m.; classes significant by one-sample t-test vs 0 are starred. Phosphatidylcholine (PC), phosphatidylethanolamine (PE), phosphatidylinositol (PI), sphingomyelin (SM), ceramide (Cer) and cardiolipin (CL) increase, whereas cholesteryl esters (CE) decrease. (B) Very-long-chain poly-unsaturated phospholipids that are selectively decreased despite the overall increase in PC/PE: log2 fold-change per species, with PC 44:6, PC 44:10 and PE 40:5 highlighted (raw p < 0.05). (C) Cardiolipins (CL) are elevated, with the linoleoyl-rich species CL(18:2_18:2_18:2_18:2) highlighted (raw p < 0.05), consistent with mitochondrial membrane adaptation to oxidative stress. (D–G) Per-class chain-length analysis: significant lipid species plotted as total acyl-chain carbons (x) against log2 fold-change (sgJMJD5/Scramble; y), coloured by degree of unsaturation (total double bonds) and sized by significance, for (D) ceramides (Cer), (E) sphingomyelins (SM), (F) cholesteryl esters (CE) and (G) triacylglycerols (TG). CEs are uniformly decreased; TGs show a chain-length switch (shorter-chain species increased, longer-chain species decreased).

