## Supplementary figures and images for "JMJD5 regulates metabolism by hydroxylating ISY1, and regulating the Arginine Methyltransferase PRMT6"

### Extended Data Figure 1

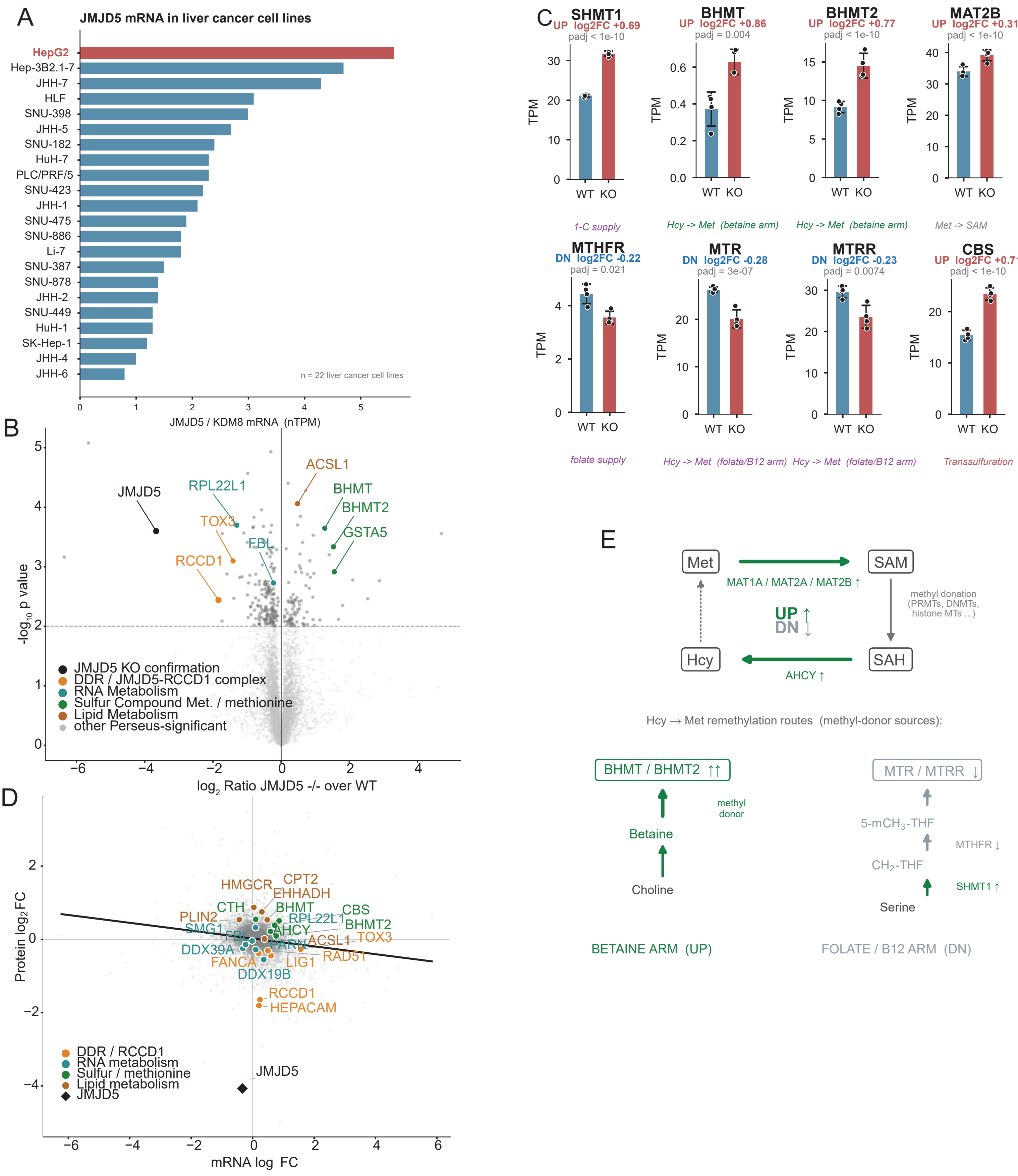

### Extended Data Figure 2

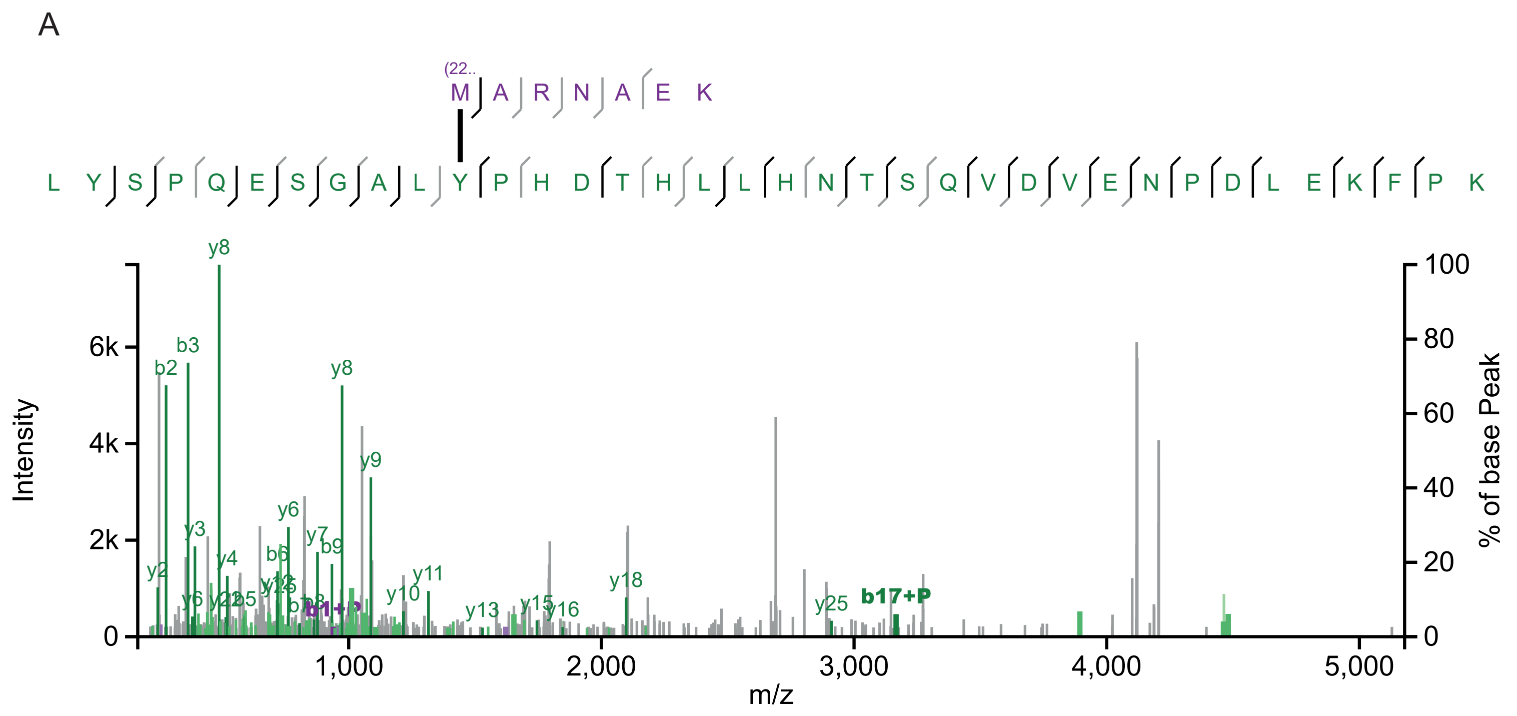

### Extended Data Figure 3

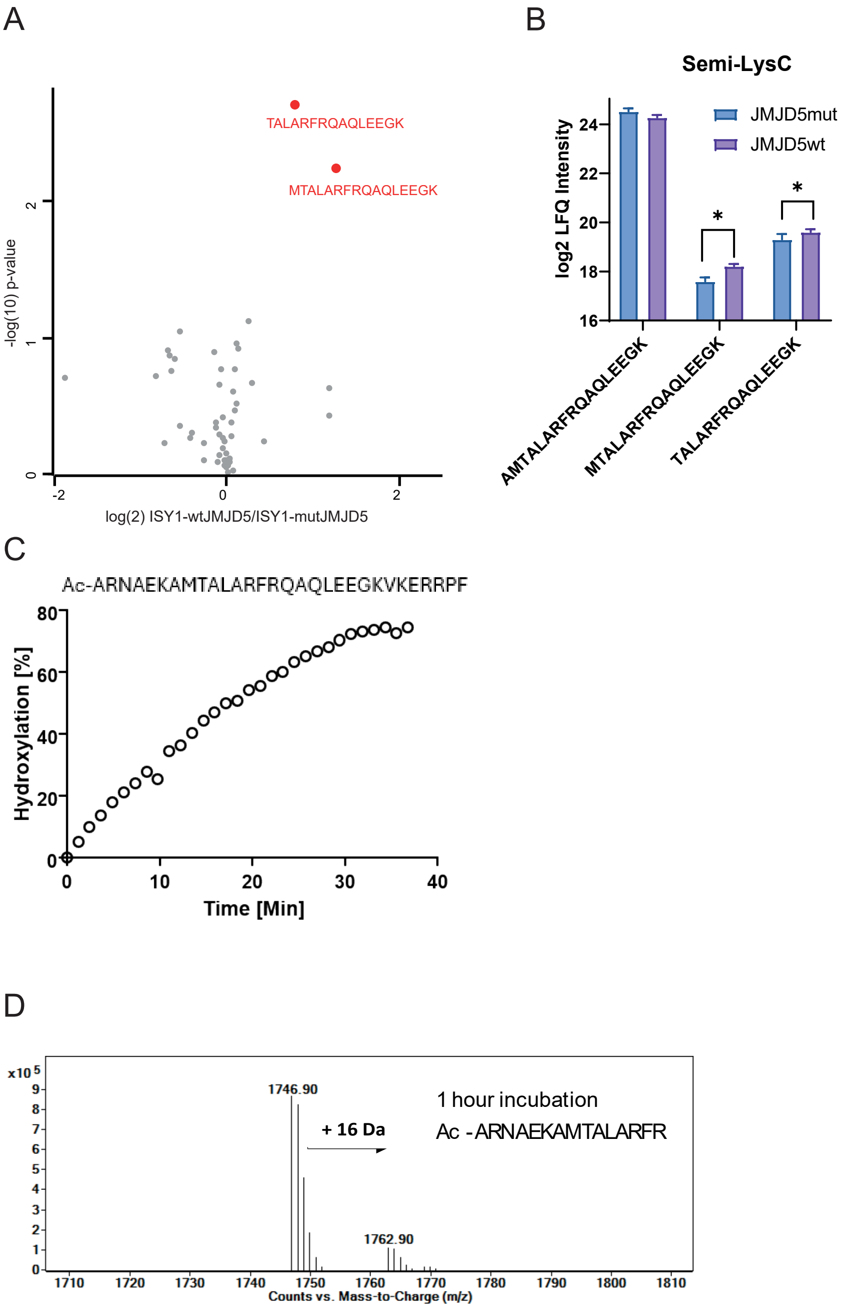

### Extended Data Figure 4

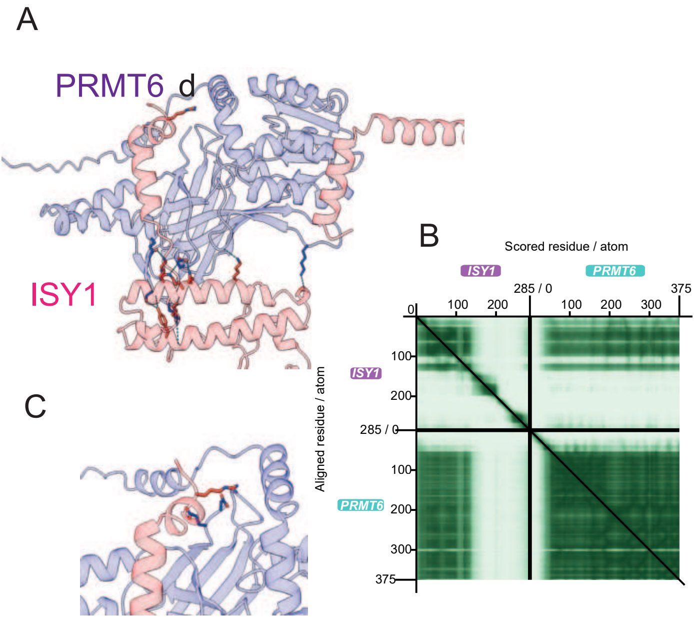

### Extended Data Figure 5

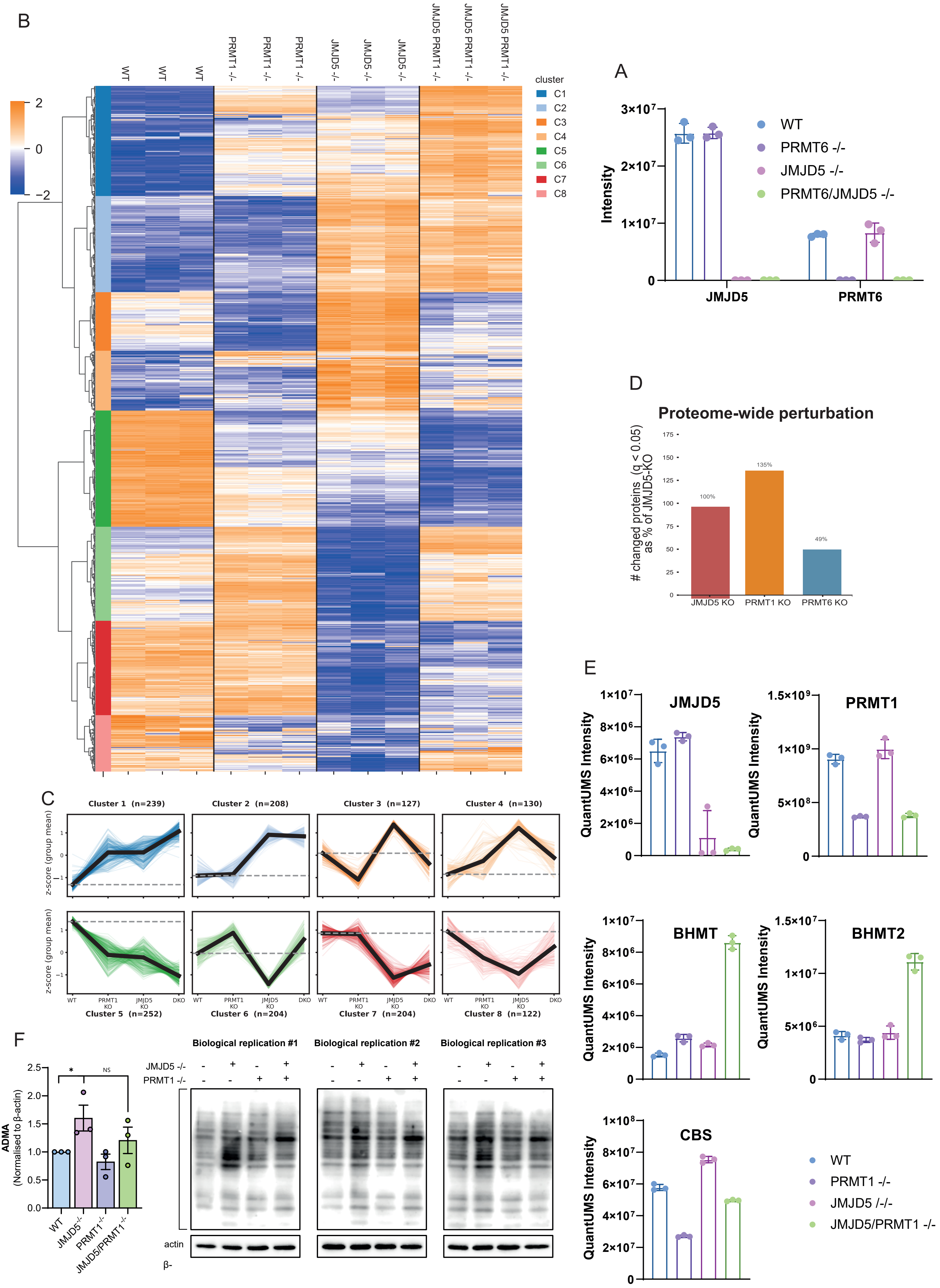

### Extended Data Figure 7

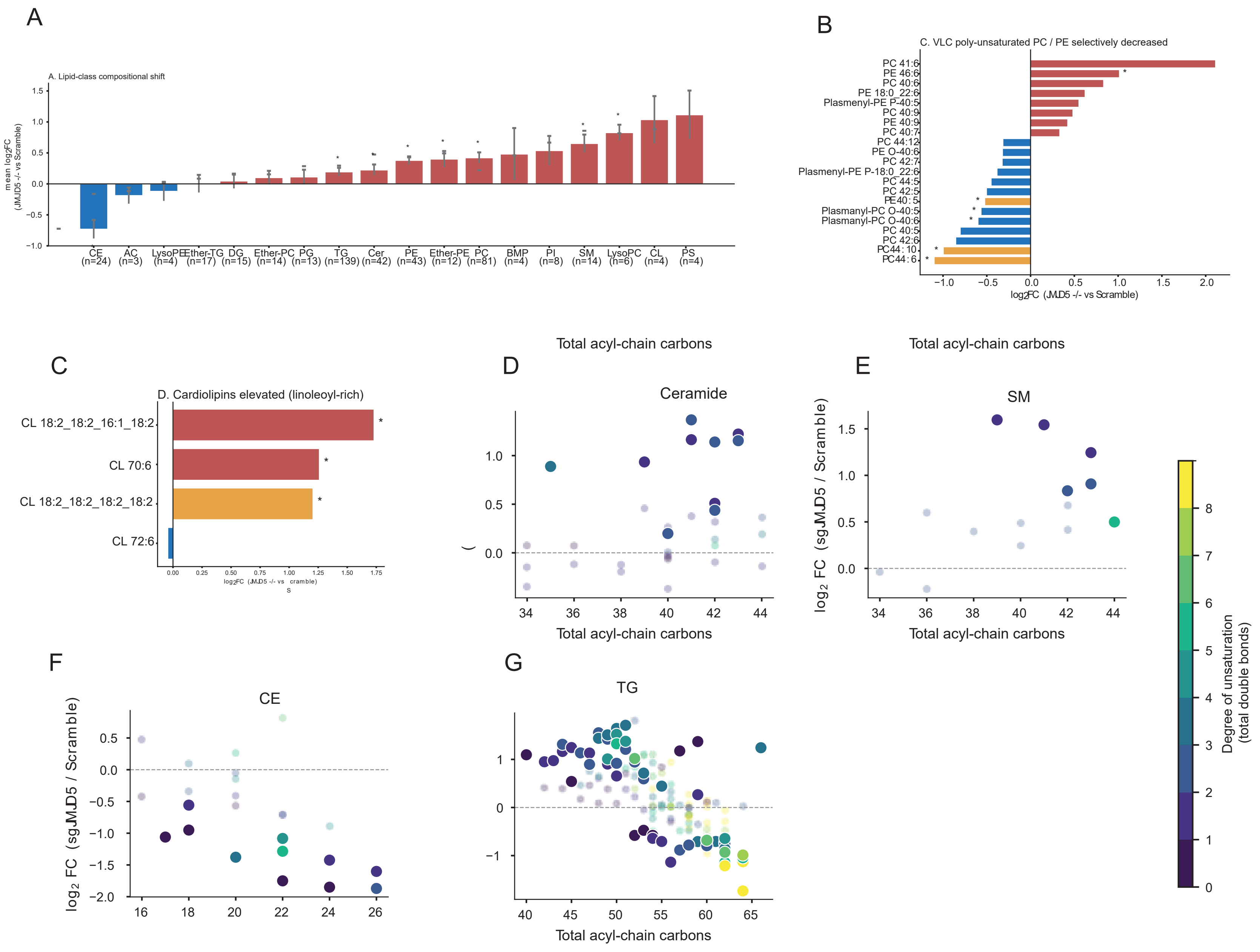
