## Extended Data Figure 6 for "JMJD5 regulates metabolism by hydroxylating ISY1, and regulating the Arginine Methyltransferase PRMT6"

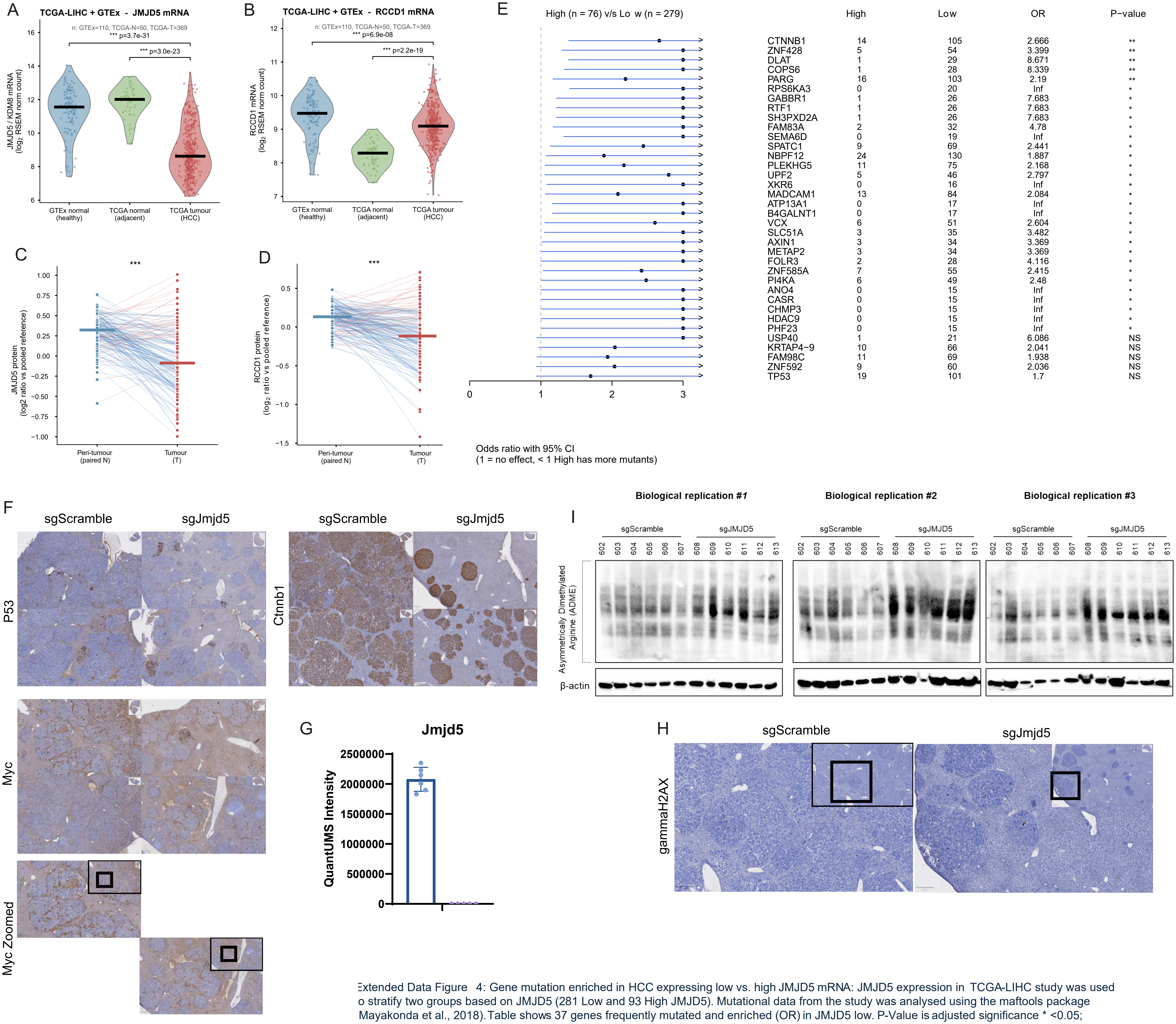


Extended Data Figure 4: Gene mutation enriched in HCC expressing low vs. high JMJD5 mRNA: JMJD5 expression in TCGA-LIHC study was used o stratify two groups based on JMJD5 (281 Low and 93 High JMJD5). Mutational data from the study was analysed using the maftools package Mayakonda et al., 2018). Table shows 37 genes frequently mutated and enriched (OR) in JMJD5 low. P-Value is adjusted significance * <0.05; *<0.01
